# Contrastive learning unites sequence and structure in a global representation of protein space

**DOI:** 10.1101/2025.09.05.674454

**Authors:** Guy Yanai, Gabriel Axel, Liam M. Longo, Nir Ben-Tal, Rachel Kolodny

## Abstract

Amino acid sequence dictates the three-dimensional structure and biological function of proteins. Yet, despite decades of research, our understanding of the interplay between sequence and structure is incomplete. To meet this challenge, we introduce Contrastive Learning Sequence-Structure (CLSS), an AI-based contrastive learning model trained to co-embed sequence and structure information in a self-supervised manner. We trained CLSS on large and diverse sets of protein building blocks called domains. CLSS represents both sequences and structures as vectors in the same high-dimensional space, where distance relates to sequence-structure similarity. Thus, CLSS provides a natural way to represent the protein universe, reflecting evolutionary relationships, as well as structural changes. We find that CLSS refines expert knowledge about the global organization of protein space, and highlights transitional forms that resist hierarchical classification. CLSS reveals linkage between domains of seemingly separate lineages, thereby significantly improving our understanding of evolutionary design.

## Introduction

Mapping the protein universe – that is, constructing representations for the relationships between all proteins – is essential to understanding the forces that have shaped protein emergence and evolution over the past 4 billion years [1-3]. To date, the most significant mapping achievements are hierarchical classifications [4-7], where building blocks of protein tertiary structure, called domains, are grouped based on sequence and/or structure similarity. Constructed with expert supervision, these databases are a repository of human insight and intuition. Yet, hierarchical classifications permit only restricted notions of relatedness and similarity, as inter-domain relationships are coarse grained into just a few nested tiers of full domains.

To capture more subtle inter-domain relationships, networks [3,8-10] and Euclidean maps of fixed-size vector representations [11-24] have been employed. Derived from pair-wise comparisons, networks can represent cross-fold similarities, where domains from ostensibly different evolutionary lineages share common sub-domain sized segments [25-29]. Similar sequence segments present in different domains [25], including ones that bridge different folds [26-28], hint at early evolutionary events [28,30]. These shared segments highlight fragment repurposing and recontextualization in the discovery and functional evolution of folds. Euclidian maps, on the other hand, are now generated by Protein Language Models (PLMs) – state-of-the-art computational models that are trained by self-supervision from vast amounts of data. Although both network- and PLM-based representations of protein space can combine sequence and structure modalities in various forms [19-21,23,31-33], they fail to provide an intuitive, joint representation of sequence and structure information.

A meaningful overview of the protein universe must provide a unified account of sequence and structure similarity, while accommodating the many-to-many relationship between them [34-36]. Moreover, by co-embedding multiple modalities, the paired data can guide each other, leading to enhanced performance. Contrastive Language-Image Pre-training (CLIP) [37], for example, co-embeds images and their natural language text descriptions. In CLIP, contrastive loss (CL) guides the image-text co-embedding, enabling state-of-the-art classification of images based on textual descriptions. We reason that the application of CL, and its pairing with protein language models (PLMs) [17,18], presents a unique opportunity to explicitly consider the complex interplay between sequence and structure.

Here, we report *Contrastive Learning Sequence-Structure* (CLSS), a protein language model, that embeds sequence segments using their corresponding structures as a guide, thereby co-embedding protein sequence and structure information. Inspired by CLIP, we used self-supervised contrastive learning on sequence and structure embeddings. Encouragingly, our trained model yields excellent agreement with curated labels including fine details, for ∼30,000 representative protein domains. Crucially, curated labels were not used during the training protocol. In cases where seemingly disparate domains have similar embeddings, visual inspection reveals that CLSS embeddings are more reflective of structural similarities than expert-curated hierarchical classification. Our trained model provides meaningful maps of protein sequence and structure, and the relationships between them, offering an unprecedented, holistic view of the protein universe.

## Results

### The CLSS architecture

We designed and trained CLSS to co-embed protein sequence segments alongside domain structures (**Figure 1;** see **Methods** for more details). Briefly, CLSS is a deep network with two towers [37,38], one for structure and one for sequence. The structure tower is based on the frozen ESM3 structure encoder, and the sequence tower has the same architecture and initial weights as the ESM2 35-million-parameter model. Both encoders are followed by a trained adapter that averages the embeddings, reduces their size, and normalizes the output. Training over 80 epochs, we minimized contrastive loss in batches of 1440 domains culled from the domains predicted by AlphaFold 2 (AF2) [39]. Full domain structures were trained against randomly chosen sub-segments of varying length, from 10 residues to full domain, with the goal of encoding contextual information within domain segments. Importantly, training is self-supervised, using only the sequences and structures of the domains.

**Figure 1.**
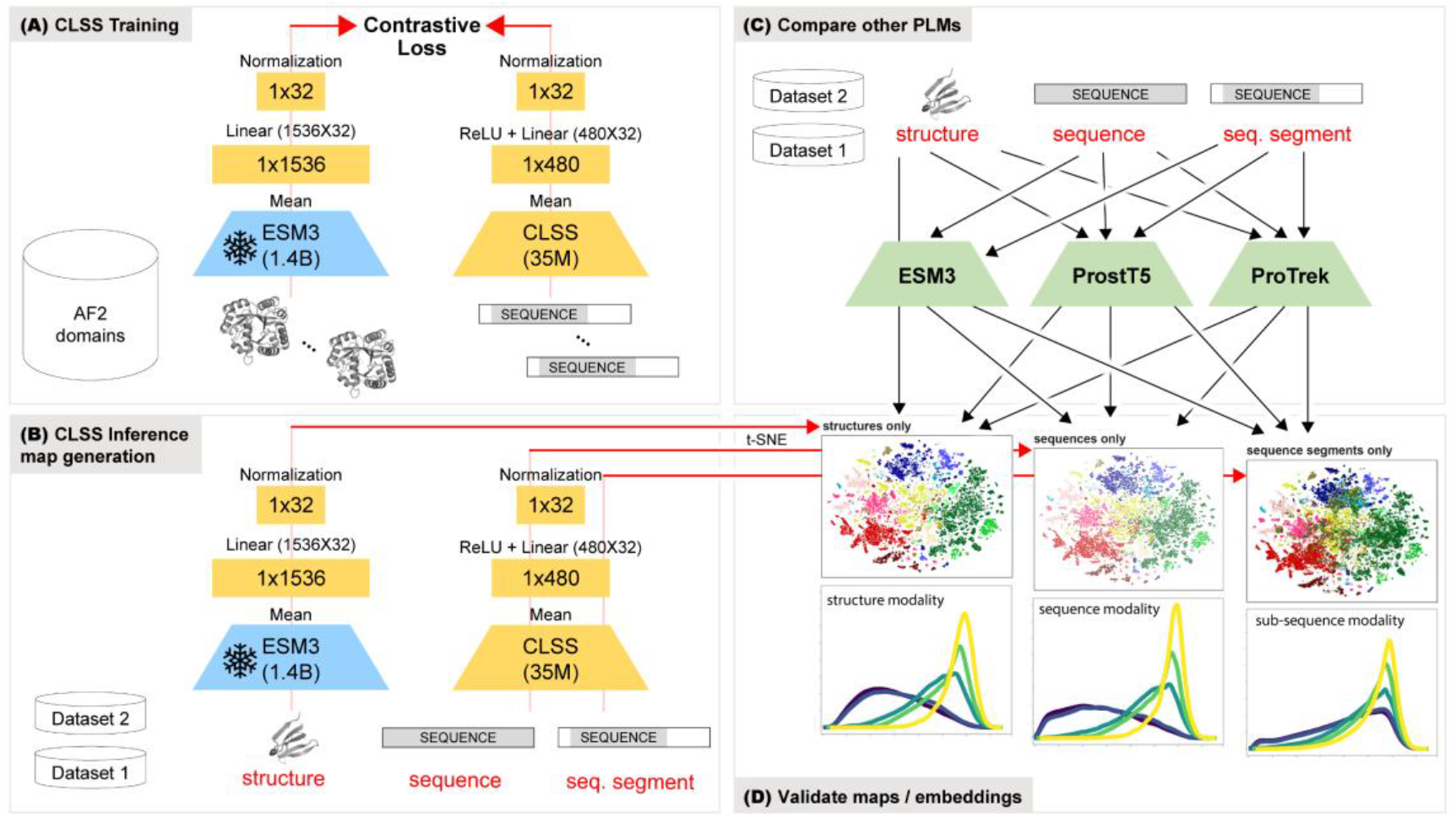
Overview of training, validating, and testing of CLSS to create unified maps of protein sequence and structure space. (A) Overview of the two-tower CLSS architecture. On the left is a structure tower based on the frozen, pre-trained ESM3 model (light blue) followed by a trained (yellow) adapter that averages, reduces dimension, and normalizes the embedding. On the right is the trained CLSS sequence tower, build upon a pre-trained ESM2 model, and its adapter network (yellow). The networks are trained using contrastive loss on batches of randomly chosen structures and sequence segments from the ECOD-AF2 domain database. Labels from a hierarchical classification were not using during training in any way. Once trained, we calculate the embeddings of the structures, sequences, and sequence segments from Datasets 1 and 2 using CLSS (B) and other PLMs (C). (D) Dimensionality reduction by t-SNE was used to create visual maps of protein space (upper images). Pairwise distance distributions (lower images) were calculated from the embeddings directly, rather than the t-SNE reduced space.

### CLSS unifies sequence space and structure space

Using the trained model, we calculated sequence and structure embeddings for two sets of representative domains taken from across the protein universe: *Dataset 1* comprises 31,696 ECOD domains taken from the 109 best-characterized protein evolutionary lineages (ECOD X-group), spanning 16 architectures. *Dataset 2* comprises 9,899 CATH domains from Røgen’s set of sequence-similar but topologically distinct domain pairs [35]. For each domain, we calculated embeddings for three modalities: structure, full sequence, and a randomly selected sequence segment of length 20-60 residues (95,088 and 29,697 embeddings for Datasets 1 and 2, respectively). We visualized the embeddings as maps: two-dimensional t-SNE projections [40] of CLSS embeddings, where each point represents the embedding of one modality of a single domain.

Maps of Dataset 1 (**Figure 2**) show that, regardless of the data modality, CLSS embeddings agree well with structure labels from ECOD, placing domains from the same architecture, and even the same structure class (*i*.*e*., all-α, all-β, α/β, and α+β domains), close to each other. Most importantly, maps from structure and sequence inputs are nearly identical, demonstrating that the model successfully co-embeds these two modalities. The distance distributions between the 32-dimensional CLSS embeddings from different modalities of the same domain in Dataset 1 (**Figure 3A**) show that the embeddings of sequence and structure are close to each other. This is true for all ECOD architectures, as seen in the architecture-dependent cumulative distance distributions between modalities (**Figure S6**). For comparison, embeddings from three other state-of-the-art PLMs – ESM3 (mean of the per-residue embeddings) [19], ProstT5 (mean of per-residue embeddings) [41], and ProTrek (the first embedded vector) [21], were calculated (**Figure 3, Figures S2, S3, S4**) – and found in separate regions of the joint map, rather than overlapping as in CLSS.

**Figure 2.**
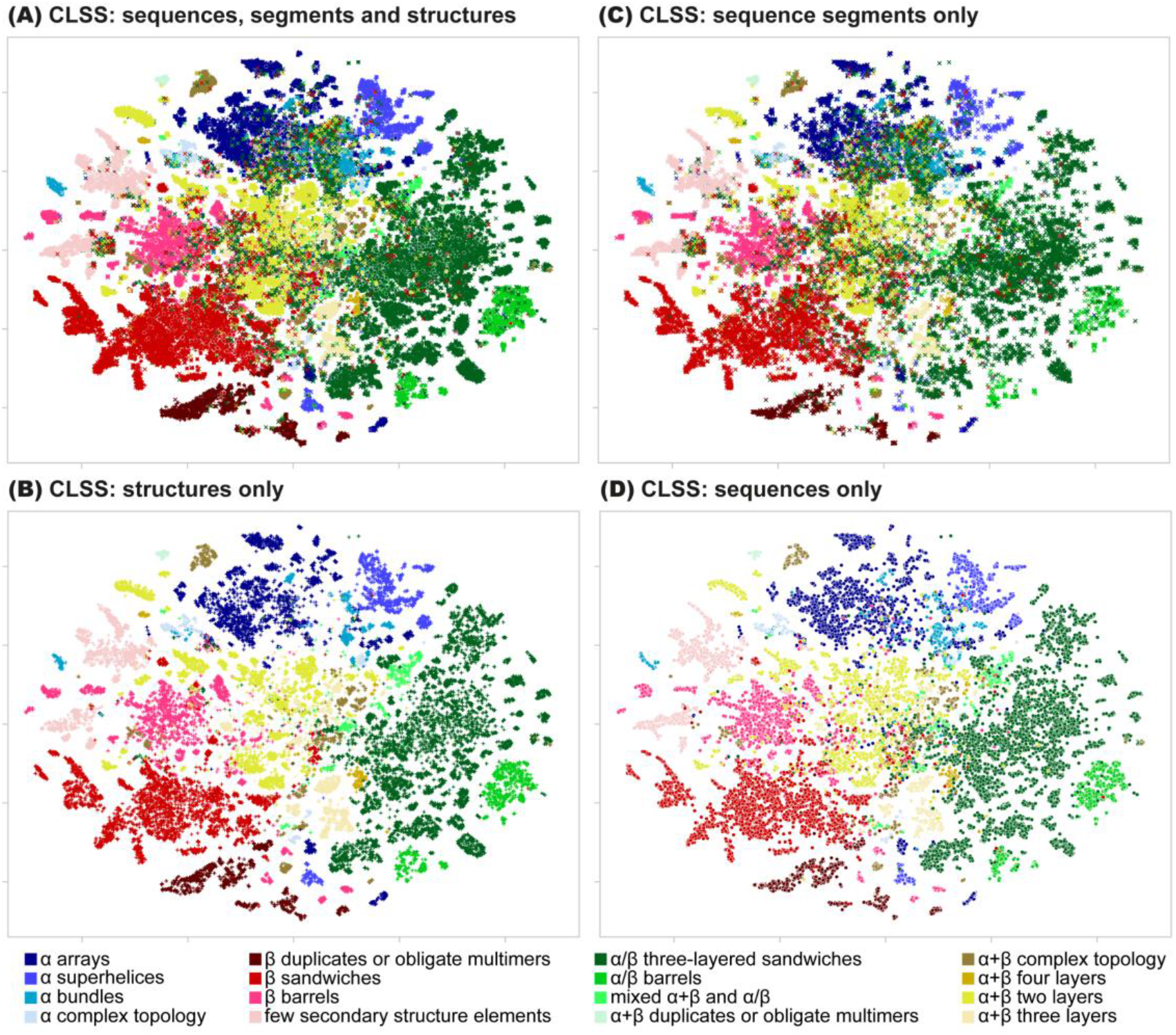
CLSS embedding maps of ECOD domains (Dataset 1). For each domain, we calculate the embeddings by three modalities – structure, sequence, and a random sequence segment – and then compute a t-SNE projection of the embeddings. Each point represents one of the modalities of a domain colored according to the label of its ECOD architecture. (A) An overlay of all three modalities. Sequences are marked by circles, structures by ‘+’, and random sequence segments by ‘x’. (B) Structure embeddings. (C) sequence segment embeddings. (D) Sequence embeddings. We find that the maps of all three modalities are very similar to each other, with the sequence (D) and structure (B) embeddings being the closest. This shows that CLSS successfully injected structure information into the sequence modality. The global organization of the CLSS embedding space positions domains with the same ECOD architecture, and even the same structure class, near each other.

**Figure 3.**
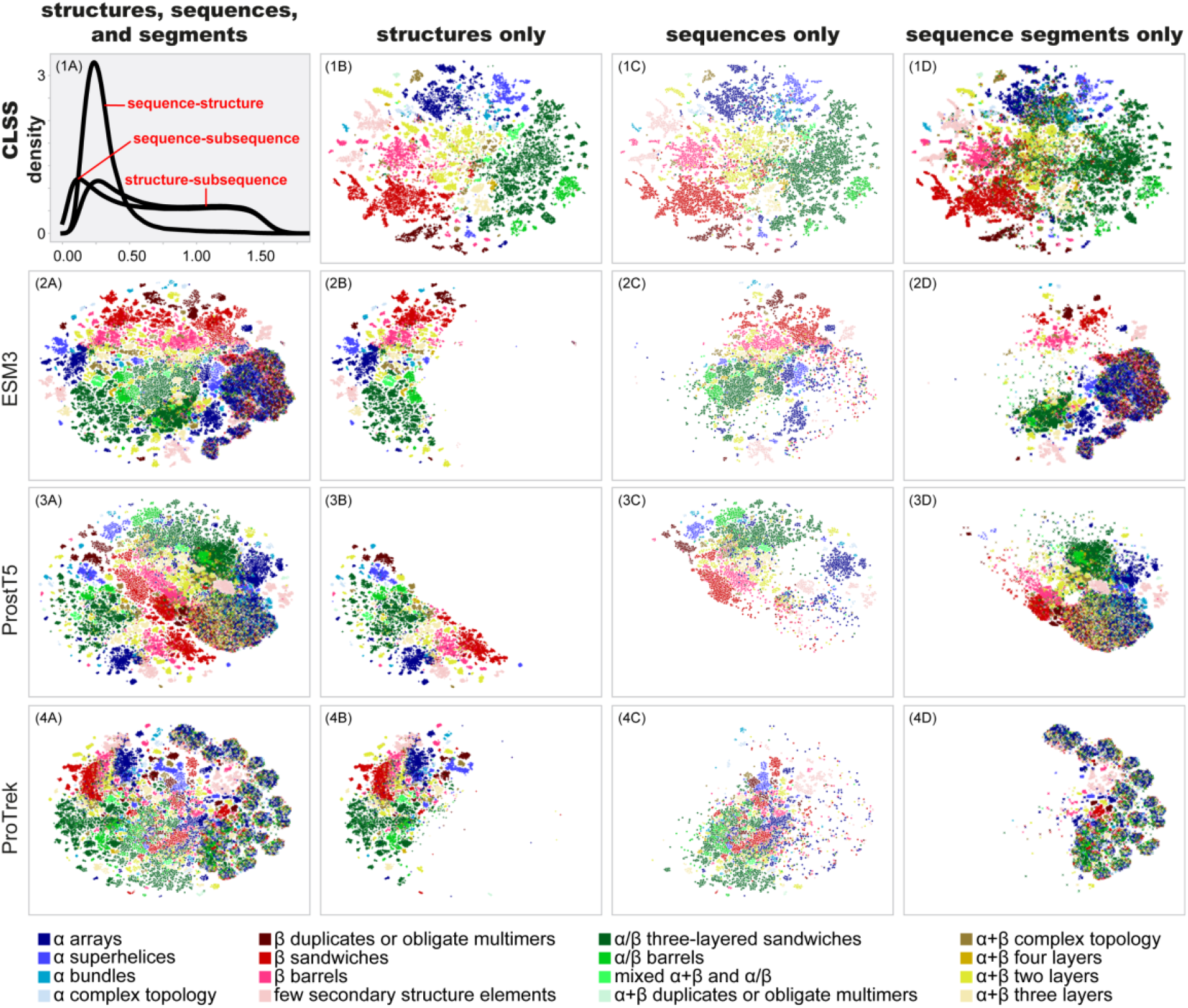
Comparison of maps generated for embeddings of ECOD domains (Dataset 1) with the PLMs (from top to bottom): CLSS, ESM3, ProstT5, and ProTrek. In each row, the maps (left to right) are an overlay of all three modalities, followed by the single modalities of structure, sequence, and sequence segments. The top-left panel shows the distribution of distances between CLSS embeddings of the same domain, in different modalities, demonstrating that the sequence and structure embeddings are indeed near each other; **Figure S6** shows a breakdown of this data by ECOD architecture. We find that in ESM3, ProstT5, and ProTrek, the embeddings of different modalities occupy distinct regions of the map. The structure modality embeddings are closer to the sequence modality embeddings, which in turn are closer to the embeddings of the sub-sequence modality. Embeddings from the structure modality generally place domains of the same architecture, and even structure class close to each other; the sequence modality performs less well (see supplementary **Figures S2-S4** for results for each PLM).

### CLSS recapitulates expert-curated hierarchical classifications

To assess whether expert-curated hierarchical classifications are reflected in the CLSS embeddings, pairwise distances were calculated for Dataset 1 domains of varying degrees of similarity. Distributions were then estimated using a Gaussian kernel density function. Both structure (**Figure 4A**) and sequence (**Figure 4C**) modalities yield virtually identical distributions, consistent with their highly correlated embeddings (**Figure 2** and **Figure 3, panel 1A**). Even though hierarchical labels were never used during training (as in some other frameworks [42,43]), we find that embedding distance distributions follow the ECOD hierarchy, where domains from successively broader groupings have more and more dissimilar embeddings. Recall that ECOD H-groups, and to some extent X-groups, attempt to capture homologous relationships between domains, while architecture and class (all-α, all-β, α/β, and α+β) are structural classifications only. Nevertheless, both structure and sequence modalities separate the distributions of all levels. In contrast, the embedding distance distributions from other PLMs is significantly less informative of sequence or structure similarity (**Figure S4**).

**Figure 4.**
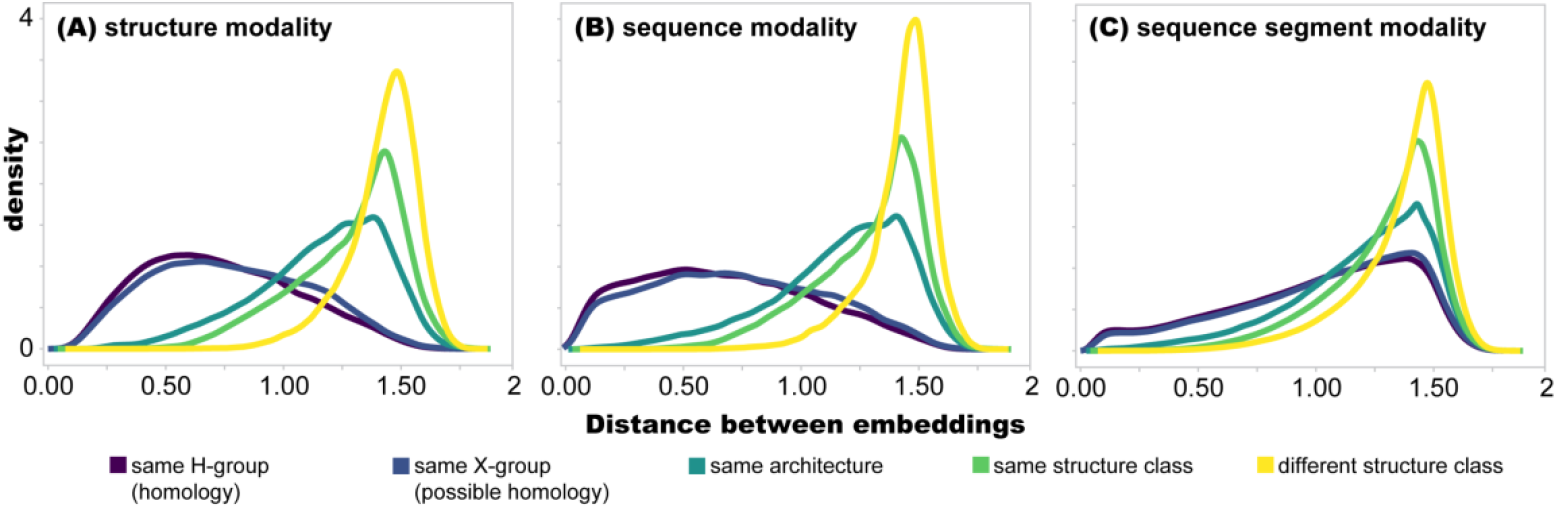
Estimated distributions of distances between pairs of embeddings for domains in Dataset 1. (A-C) The distributions of distances of domain pairs with the same ECOD label at different levels of the hierarchy. The color palette indicates the shared level in the ECOD hierarchy, from pairs with the same H-group label in purple through pairs with different structure class in yellow. In all three modalities, the distances follow the ECOD hierarchy, with the domain pairs from the same H-group being closest. Thus, CLSS embeddings for all three modalities agree with the human-curated labels of ECOD. Notice that the distance distributions of the H- and X-group pairs for the sequence modality (panel B) are slightly shifted to the left compared to the structure modality (panel A), implying that there are pairs within these groups whose sequence embeddings are closer to each other than their structure embeddings.

### CLSS robustly co-embeds the sequence and structure of metamorphic domains

Co-embedding of proteins with similar sequences but dissimilar structures poses a unique challenge to contrastive learning: In the sequence modality, the embeddings should be near each other (due to sequence similarity) but in the structure modality the embeddings should be far apart (due to structural dissimilarity). To understand if and how CLSS resolves these conflicting signals, we mapped the ‘metamorphic’ CATH domains of Dataset 2. To our surprise, the sequence and structure maps are generally similar to each other (**Figure 5, Figure S10, panel A**) and both modalities successfully group domains from the same architecture and super-architecture (**Figures 5A, 5B** and **5D**). As before, CATH labels were never used during training. This result is likely a consequence of training on sequence segments, as diverse segments can be extracted from a fold but CLSS still places these segments near the full domain in embedding space, described in detail below. Although the other PLMs compared here do not co-embed modalities – and thus, one does not expect metamorphic proteins to present a challenge – the resulting groupings are far less pronounced (**Figures S7, S8, S9**; *e*.*g*., α-domains). Finally, a joint map of domains from Dataset 1 and Dataset 2 shows that the metamorphic CATH domains fall near their corresponding ECOD X-groups (**Figure S10**; note that t-SNE projections depend on the dataset mapped such that by including the diverse domains of Dataset 1 in the map, some CATH domain clusters contract).

**Figure 5.**
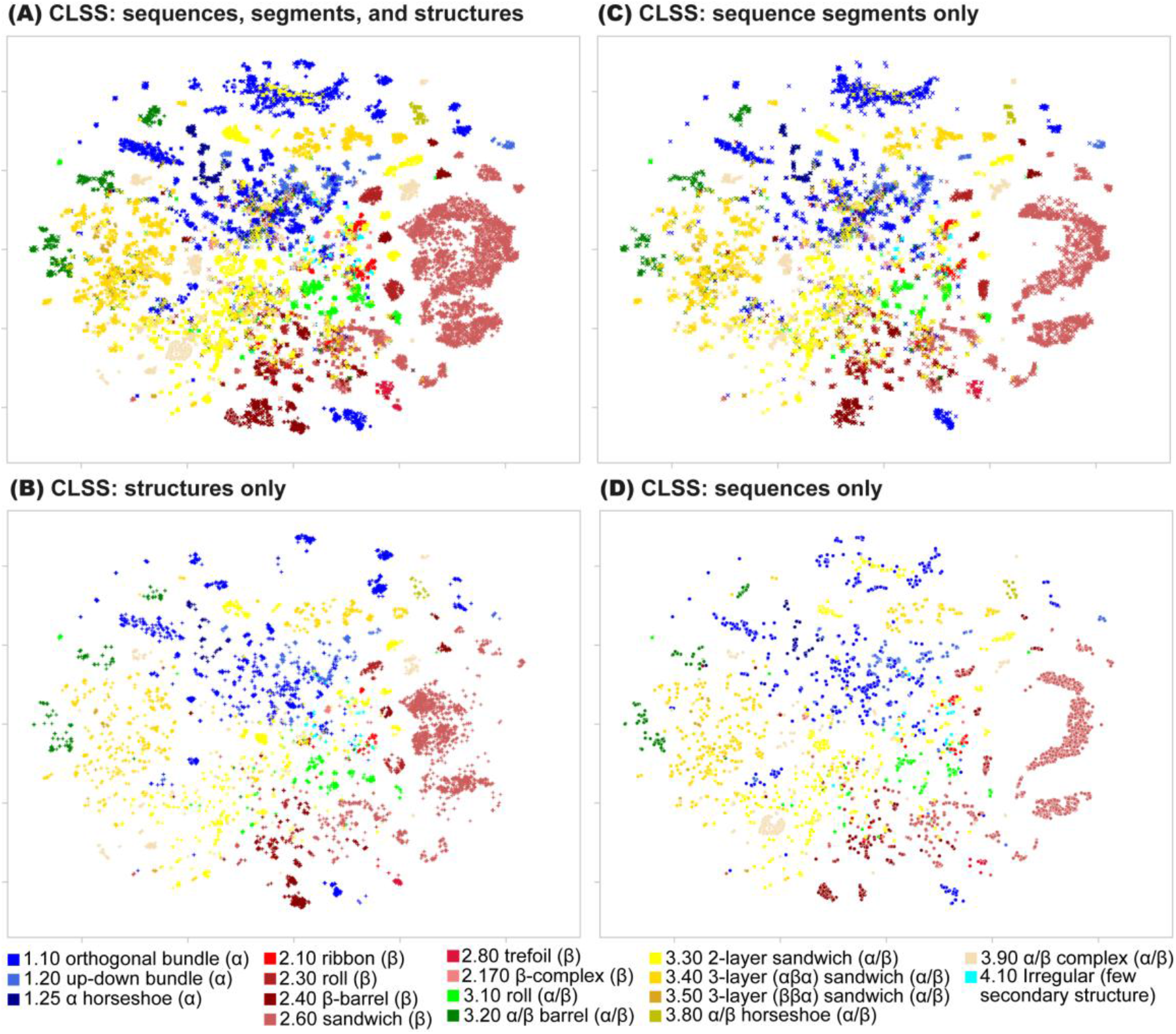
Maps based on CLSS embeddings of metamorphic CATH domains, sharing high sequence similarity but topologically different structures (Dataset 2). For each domain, we consider three modalities: sequence, structure, and a random sequence segment and compute a t-SNE projection of the embeddings. Each point represents one of the modalities of one domain colored according to the label of its CATH architecture. (A) All three modalities. Sequences are marked by circles, structures by ‘+’, and random sequence segments by ‘x’. (B) Structure embeddings. (C) sequence segment embeddings. (D) Sequence embeddings.

### CLSS encodes the context of domain sub-sequences

CLSS training included sequence segments of at least 10 residues. Thus, CLSS should be able to accurately embed domain sub-sequences as a third modality. The maps and distributions in **Figures 1-5** and **S2-S5** include embeddings for the sub-sequence modality. **Figure 2C** shows the map of the sub-sequence CLSS embeddings for Dataset 1, which is broadly similar to the maps obtained using the full sequence (**Figure 2D**) and the full structure (**Figure 2B**). Moreover, the embedding distance distributions from the sub-sequence modality (**Figure 4C**) follow the same trend as the other modalities, with domains from broader groupings resulting in progressively more distant embeddings. Nevertheless, embedding distance distributions for H-group and X-group level similarity are further away from each other than for the other two modalities. The distance distributions between embeddings from different modalities of the same domain in Dataset 1 (**Figure 2, panel 1A** and **Figure S4**) show that sub-sequence and sequence yield more similar embeddings than sub-sequence and structure. Neither ESM3, ProstT5, nor ProTrek yield informative embeddings for the sub-sequence modality (**Figures 3D** and **S4**); their embedding distance distributions are virtually the same regardless of the level of the shared labels.

There are cases where the embedding of a sub-sequence is far from the embeddings of the full domain from which it came, even for CLSS. Often, these cases are associated with architectures that contain both α and β elements: Primarily mixed α+β and α/β (light green), α/β three-layered-sandwich (dark green), α/β barrel (green), and, to a lesser extent, α+β complex topology (brown) and α+β four layers (light brown; discussed in more detail below). This may reflect that α/β folds have significant fragment similarity between them, such that extracting a short fragment greatly diminishes information about the surrounding domain context.

### Comparisons between CLSS maps and hierarchical classifications can highlight transitional proteins

Are differences between hierarchical classifications and CLSS maps due to the limitations of CLSS, or do they reveal *bona fide* domain features that are not well-captured by hierarchical classification? To explore this question, **Figure 6** shows two instructive comparisons between ECOD and CLSS; a third example is described in the SI and **Figure S16**.

**Figure 6.**
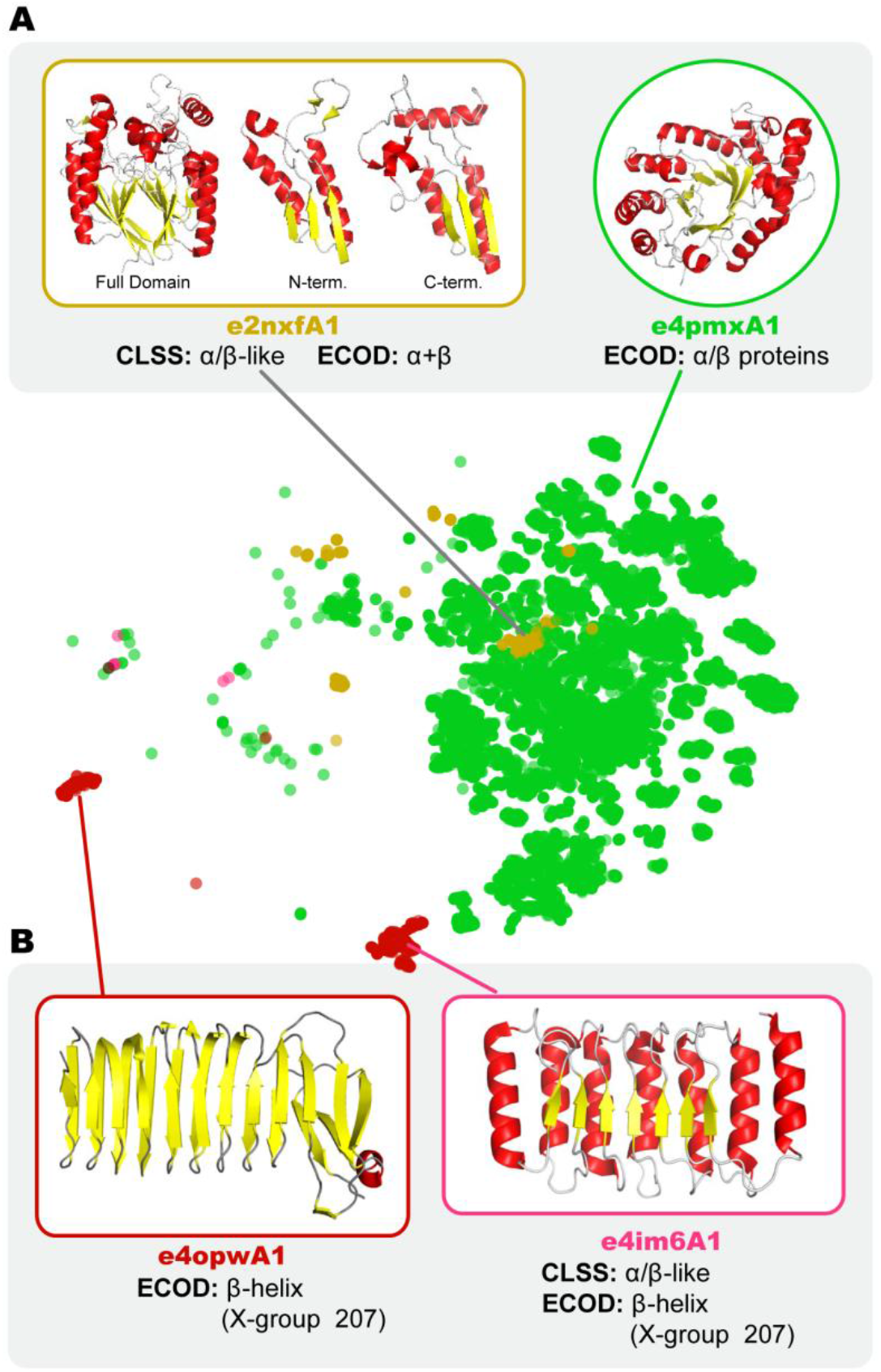
Example comparisons between the ECOD hierarchy and CLSS maps. (A) The localization of some α+β 4-layer sandwiches (ochre) with α/β-folds (green). The CLSS classification reflects the β-α-β-α-β elements that are contained within their structures. (C) The splitting of X-group 207, which is classified by ECOD as having a β-architecture. Domains positioned near the α/β region (green) have clear signatures of α/β-structure. These examples, and another described in the supplementary text (Figure S16), highlight the limitations of hierarchical classifications and present three candidate events of structural evolution within and between protein X-groups.

One example of a difference between the two approaches is the localization of α+β four-layer sandwiches within the greater α/β cluster on the CLSS map (**Figure 6A**; marked in ochre). Inspection of the structure, however, affirms the CLSS clustering by revealing significant α/β-character. Domain e2nxfA1 (X-group 246), for example, has two juxtaposed β-α-β-α-β motifs that form the core of the fold. However, unlike a Rossmann domain – in which the two β-α-β-α-β motifs form a single extended β-sheet with a “doubly wound” 3-2-1-4-5-6 strand order – the β-α-β-α-β motifs of this domain pack against each other, face-to-face, to form the four-layer α-β-β-α sandwich structure [28]. The localization of the α+β four-layer sandwiches within the larger embedding space occupied by the Rossmannoid folds suggests the evolutionary potential for β-α-β-α-β motifs to interconvert between the two possible packing arrangements, albeit with a clear preference for β-strand edge-to-edge packing.

A second example of a difference between CLSS embeddings and the ECOD hierarchy relates to the classification of single-stranded right-handed β-helices (X-group 207), hereafter referred to as β-helices. Although β-helices are classified as β proteins by ECOD, β-helix domains belonging to H-group 207.1 are located near α/β proteins on the CLSS map (green colored embeddings), whereas β-helix domains belonging to H-group 207.2 are located near other β-proteins, as would be expected (**Figure 6B**). Inspection of the structures once again affirms the CLSS embeddings: 207.1 β-helices do, in fact, have significant α/β-character. The ECOD hierarchical classifications scheme permits only a single architecture classification per grouping, and the β-helix highlights how this limitation can be misleading for groups with diverse structures. If H-groups 207.1 and 207.2 are indeed related, as ECOD suggests, the β-helices represent a remarkable case of protein structure exploration, bridging the space of β-folds with that of α/β-folds.

In short, differences between hierarchical classification schemes, represented here by the ECOD database, and CLSS seem to reflect the rigidity inherent to hierarchical classification. Moreover, these examples suggest that protein space is more continuous than hierarchal in nature.

### Cofactor utilization approximately bisects the protein universe

Coloring the CLSS map based on protein structure class, cofactor, and metal associations (**Figure 7**) reveals a striking split between domains with few cofactor associations (bottom-left side of panel C) and those with strong cofactor associations (upper-right side of panel C). Comparing **Figures 7A** and 7**C**, we find that cofactor association is very common among α/β and α+β domains, but much less common in α-rich domains, and very rare in β-rich domains. While the prevalence of α/β folds in metabolism has been noted previously [1-3], the map view afforded by CLSS brings this functional preference into sharp focus. It is tantalizing to hypothesize that the structure features of α/β and α+β folds, which often have α/β elements, confer a fundamental preference for cofactor binding. The special properties of βαβ fragments, and particularly of the N-terminus of the α-helix and the loop that precedes it [5-7], may be the causal factor, though historical explanations cannot be ruled out.

**Figure 7.**
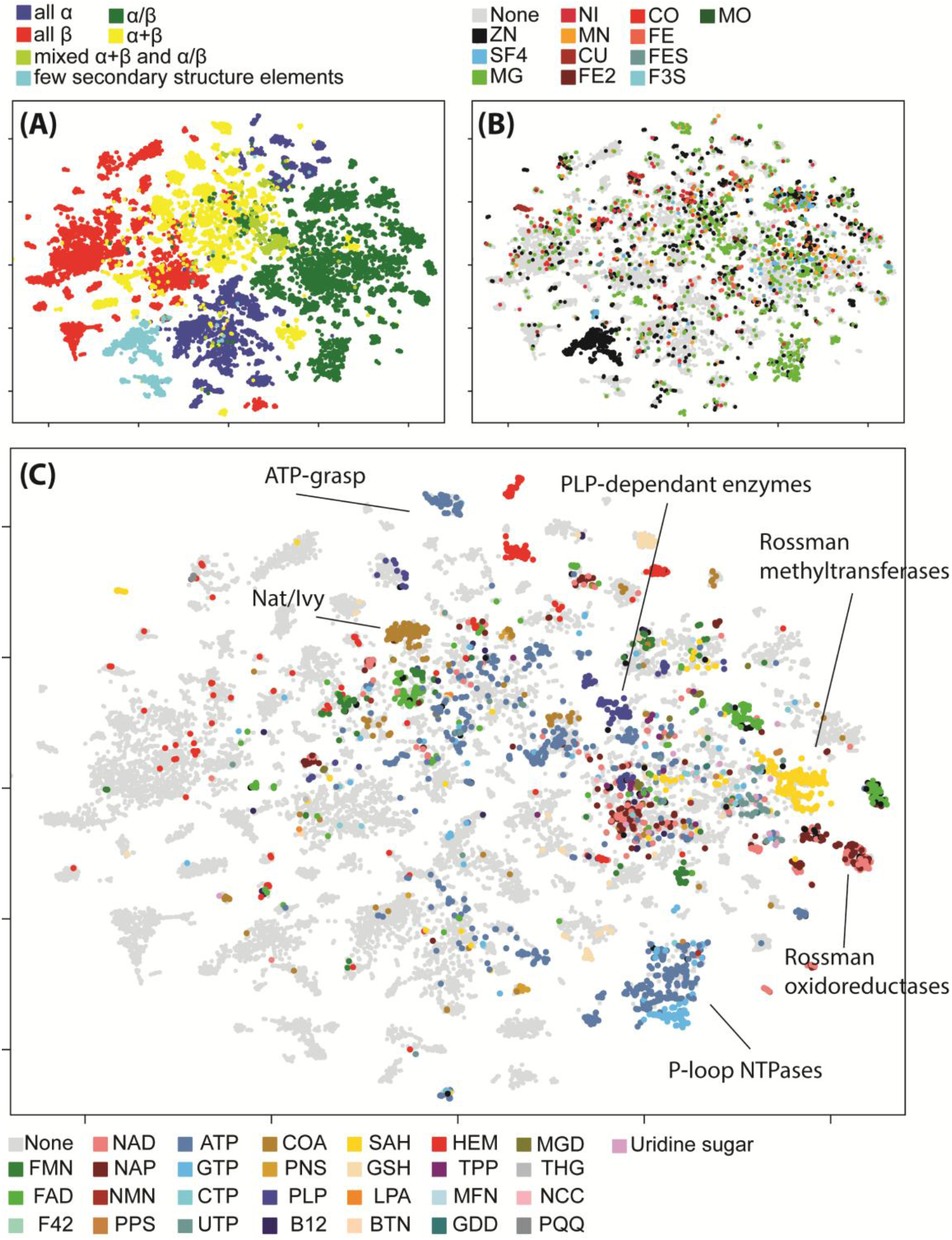
Cofactor utilization approximately bisects the protein universe. A map of the CLSS sequence embeddings of the ECOD dataset (Figure 2D), colored by: (A) ECOD class, (B) metal binding association, and (C) organic cofactors association. Metal association is found throughout. However, there is an enrichment in cofactor association in the upper right region (corresponding to α/β and α+β ECOD classes) compared to the lower left region (corresponding to all α and all β classes).

The pattern of organic cofactor associations is unlike that of metals (**Figure 7B**), which is more sparsely distributed across almost all regions of the map. A few regions, however, have significant enrichment for a specific metal association. For example, zinc cations are strongly associated with a cluster of several X-groups annotated as having the “few secondary structure elements” architecture. A second cluster of domains from the same architecture, but that do not have strong associations with zinc, is close by, suggesting that CLSS near-hierarchically grouped these two clusters together (owing to their broad lack of secondary structure) and then splits them according to the structure and sequence features associated with zinc binding. Other strong metal associations are in full agreement with expectation: Magnesium cation associations are enriched in the region of P-Loop NTPases, which have a conserved Mg2+ cation in the active site; and cupredoxins are strongly associated with copper ions.

### The CLSS viewer

We provide a web interface for exploring a t-SNE projection of the CLSS embeddings of almost all ECOD domains from the ECOD-AF2 F100 dataset (https://gabiaxel.github.io/clss-viewer/). Individual domains or groups of domains can be searched for by their domain identifier (e.g., “e6tjeA1”), evolutionary grouping (e.g., “207”, “207.1”), and partial name (e.g., “lectin”). When an area of interest on the plot is selected, the list of associated domains, along with their ECOD classifications and (x, y) coordinates, are displayed. To facilitate data sharing, the URL is updated to stably reflect the current selection state. In addition, selected domain data can be downloaded as a CSV file for offline analysis.

## Discussion

We introduce CLSS, a self-supervised sequence-structure co-embedding model, and demonstrate its use in mapping the protein universe. Given a query protein sequence, or structure, or even only a sequence segment, CLSS can embed it next to similar or related domains. The CLSS viewer provides an easy-to-use, interactive interface for exploring CLSS embeddings.

CLSS differs from established PLMs in several important ways: First, CLSS was trained on sub-domain sequence segments and the structure of the full domain within which the segments are situated; CLSS is not a masked protein language model. Consequently, CLSS embeddings contain meaningful domain, and even sub-domain, representations. Second, the contrastive learning procedure successfully co-embeds sequence and structure information. Importantly, we use a CLIP-inspired self-supervised contrastive loss approach, which considers a randomly chosen large batch of proteins and contrasts the distances between the embeddings of the different modalities. With no use of hierarchical labels during training, any similarity to hierarchical classifications in inference is emergent. Our approach is starkly different from a loss function that contrasts the embeddings of a triplet selected using the labels in a hierarchy [43], which is also confusingly dubbed contrastive. Because of these two novel qualities, CLSS-derived representations of the protein universe are superior to those of ESM3, ProstT5, and ProTrek, as judged by its ability to reproduce known evolutionary relationships. CLSS also includes several quality-of-life improvements: With only 36M trainable parameters, it is considerably smaller than other state-of-the-art models, even those considered lean [44]. Thus, training purpose-built models with CLSS-like architectures is possible with only modest computational resources. Finally, CLSS uses a compact embedding space of only 32-dimensions [45], facilitating data storage and searching [46]. These design choices provide an alternative view on the future of PLMs, with a better bias-variance tradeoff that creates more efficient and compact models [47].

CLSS contrastive loss aims to co-embed sequence segments, including ones far shorter than the full domain, near the structures that host them. This property can be useful when embedding metamorphic proteins because their sequence segments must be compatible with different structures. Indeed, even though co-embedding should be challenging for metamorphic domains, CLSS successfully maps CATH metamorphic domains (Dataset 2). By characterizing the sequence segments, or motifs, that can fit within the context of a specific structure, CLSS may aid in high-throughput library-based design approaches, and thus may facilitate protein design [48-51]. The motivation for sub-segment training is similar to that of protein threading [52] and early protein structure prediction approaches that quantified the compatibility between a sequence and a (potentially much larger) structure [53-55].

We argue that hierarchical domain classifications, like ECOD, are relevant and challenging benchmarks for PLMs because they curate expert knowledge about protein evolutionary interrelationships, which is informative for functional characterization [7,56,57]. Our view is a departure from current practice, which centers more on the success of a supervised downstream task, such as the prediction of Enzyme Commission (EC) numbers, gene ontology (GO) terms [20,22,23,31-33,42], or the identity of masked residues [19]. Assessment of features that are largely defined at the protein family level provide little information about the global organization of the embedding space. Instead, by painting PLM-based maps with expert knowledge of the protein universe at various degrees of abstraction [17,19,20,22,23,32,33,58], we can assess the large-scale and fine-scale structure of the embedding space with an unprecedented degree of explainability.

The agreement of the CLSS maps with hierarchical labels is striking, especially in light of its self-supervised training, which did not have access to this information (e.g., in contrast to [42,43,59]). CLSS maps from the sequence, structure, and sub-sequence modalities capture documented similarities of sequence and structure. Cases where hierarchical classifications and CLSS appear to disagree seem to highlight potential events of structural exploration: connections between different folds and possible examples of protein universe-spanning structure divergence. Although these types of relationships are not necessarily needed for protein structure prediction *per se* [53-55], they provide clues into the nature of protein sequence-structure exploration. In short, CLSS offers a new, AI-based, perspective on the continuous, malleable nature of protein sequences and structures, a long-sought scientific challenge [38]. When paired with studies of cross-fold fragment sharing and the observation of metamorphic protein folds, our data support the view that the protein universe, despite its staggering diversity, is readily traversable over long timescales. Moreover, a map-based representation overlaid with functional properties, such as cofactor and metal binding, provides new insights into the biological processes that gave rise to the protein universe.

Looking forward, we anticipate that CLSS, or models like it, will be a valuable resource for molecular biologists. CLSS maps can provide new contextualizing information – a view of protein similarity that encompasses aspects of sequence and structure. Ultimately, a unified sequence-structure embedding space will create new possibilities for database searching, sequence alignment, protein engineering, and evolutionary trajectory analysis.

## Methods

### CLSS Architecture

**Figure S1** illustrates the architecture of CLSS. Our contrastive model has a two-tower architecture: The sequence tower is a trainable ESM2-like model with 35M parameters [26]; the structure tower is a frozen ESM3 [2] model with 1.4B parameters. Given a protein (or domain) with *N* residues, the sequence and structure towers infer a matrix of size *N* × *K*, where *K* is a model-dependent constant (480 for ESM2 and 1536 for ESM3). Taking the mean over the matrix columns for each input results in vectors of length 480 and 1536 for ESM2 and ESM3, respectively. Finally, an adapter (a single linear layer) of size *K* × 32 is appended at the end of each model, followed by L2 normalization. For every sequence-structure pair, CLSS outputs two vectors of length 32.

### Datasets

Our training/validation dataset comprises 1 million randomly sampled domains with replacement from the publicly accessible ECOD domain classification of AlphaFold 2 predicted structures [39]. CLSS uses ESM3’s `ProteinChain`API and ignores domains that have PDB files the API failed to open. Following Machine Learning best practices, we randomly split the dataset into a training set with 950,000 domains, and a validation set with the remaining 50,000 domains. Dataset 1 comprises 31,696 ECOD domains taken from the 109 most-characterized protein evolutionary lineages (ECOD X-group), spanning 16 architectures. Dataset 2 comprises 9,899 CATH domains from Røgen’s set of sequence-similar but topologically distinct domain pairs [35]. The CLSS viewer shows all domains in the ECOD-AF2 F100 dataset with a valid PDB file (approximately 1.81 million) except for 95 domains with missing or corrupt PDB files that our download pipeline failed to fix. In cases where the provided PDB files were invalid, we applied the PDBFixer python library. In cases where the provided PDB files were empty, we used the PDB file from the ECOD-PDB dataset instead and, if that file was empty as well, we used the relevant PDB file from RCSB and cut it manually according to the domain’s start and end residues as listed on ECOD-PDB-development-291.

### Training Process

In each training step, 1440 domains (illustrated with a batch of size 4) were randomly selected from the full training dataset. The sampled batch was then split into 8 minibatches of 180 domains. Each minibatch was assigned to a different GPU. From the sequence of each domain, we sampled a random subsequence (highlighted in **Figure 1** in light blue) and calculated its ESM2-like sequence-segment embedding and projection to a normalized vector of size 32 (annotated in **Figure 1** as *Se*_1_,…, *Se*_4_). We then calculated the ESM3 structure embedding for the full domain and its projection to a normalized vectors of size 32 (annotated as *St*_1_,…, *St*_4_). Our contrastive loss function maximized the dot product between the structure and sequence segment embeddings from the same domain (marked in green), while minimizing it between different domains (marked in gray). Finally, we applied the `SoftMax`function to each row of the matrix, and computed the cross-entropy loss on the resulting distribution of each row, while labeling values on the diagonal as the “correct” class. Overall, we trained the network for a fixed number of 80 epochs on 8 NVIDIA A100 (40GB) GPUs using the PyTorch Lightning framework and `DDPstrategy`for approximately 4.5 days.

The training process updated all parameters of ESM2-like model and the adapter networks of both models, while keeping the parameters of the ESM3 network frozen. This design implemented our goal of injecting structure information into the sequence model. Overall, the number of trained parameters in our network was only ∼36M, slightly more than the small ESM2 (35M) model, and dramatically less than ProstT5 (3B), ESM3 (1.4B), and ProTrek (930M).

Our evaluation was inspired by CLIP: the co-embedding was trained on the pairing of extensive data, followed by evaluation on labeled data. In CLSS, the pairing is between the sequence of segments and the structure of protein domains in large datasets. Importantly, neither ECOD nor CATH labels were used during training in any way. Consequently, analysis of the trained embeddings with respect to ECOD and CATH labels cannot suffer from data leakage of these labels. By randomly extracting the training and validation domain datasets from the same distribution of domains – as opposed to enforcing that these datasets have minimal overlap – we ensure that the learned latent space is preferentially organized by the most ubiquitous domain families. For the goal of visualizing a meaningful representation of protein space, one must train on an exhaustive dataset representing it, as do all methods compared here – ProstT5, ESM3, ProTrek, and CLSS.

### Cofactor-domain and metal-domain associations

Associations between domains and ligands, either metals or organic cofactors, were extracted from the ECOD-PDB-development-291 flat file (last column), which were calculated using a 4 Å ligand-domain distance cutoff. Common cofactors, as well as their derivatives – including reactant and product forms of the same cofactor (*e*.*g*., ATP and ADP), common derivatives (*e*.*g*., CoA and acetyl-CoA), and common non-reactive analogs (*e*.*g*., phosphoaminophosphonic acid-adenylate ester for ATP) – were grouped together (**Table S1**). For clarity, domains that bind two or more ligands of the same type (organic cofactor or metal), which is less than 5% of domains that bind a ligand, were colored for just one of them.

### Ablation Studies and Hyperparameter Search

Several ablation studies and hyperparameter searches were performed. We trained the same model architecture with adapters of varying sizes: 16, 32, and 64. We also varied the minimum length of the random sub-sequences sampled from each full domain sequence (considering 1, 10, 15, 20, 25, and 30 residues), as well as training using the full-length sequences only, without any sub-sequence sampling (note that this model was only trained for 42 epochs). In addition to testing multiple architectures, we also ablated our loss function: optimizing the temperature parameter during training (similar to CLIP), versus using a fixed temperature with multiple values (0.25, 0.5, 0.75, 1, and 1.25); calculating the loss on both the rows and columns of our similarity matrix, or just on the rows; and computing the loss globally, on entire batches, compared to locally on each GPU.

To evaluate the trained models, we used the distributions of distances between each domain sequence, structure, and sub-sequence embeddings for a validation set of 50,000 ECOD domains. We find that increasing the size of the adapter networks yields higher distances between the different modalities (**Figure S12A**), diminishing performance, while decreasing it results in a slight improvement over our chosen architecture (**Figure S12B**). **Figure S13** shows that, generally, training on shorter sequence segments results in slightly higher distances between the sequence and structure modalities, but significantly lower distances between the sequence segment modality to both the sequence and structure modalities. This holds true for all tested models, except for the model trained on segments of any length (minimum length of 1 residue) (**Figure S13 E**), which performs worse than our chosen model, which was trained on segments with 10 or more residues. This exception suggests that sequences with less than 10 residues do not contain enough information to associate them with their parent domain structures. Conversely, training on full sequences (**Figure S13F**) yields vastly lower distances between the sequence and structure modalities than our chosen architecture, but has the opposite effect with respect to the sub-sequence modality, for which distance distributions to both the sequence and structure modalities are shifted significantly towards greater values. This demonstrates the significance of training on sequence segments to the success of CLSS.

Interestingly, optimizing the temperature during training yields worse results than our chosen architecture, which uses a fixed temperature of 0.5 (**Figure S14A**). Increasing the temperature to 1 generally results in better performance, while further increasing it does not have a significant effect (**Figure S14**, panels B-E). The models trained with a fixed temperature of 1 and 1.25 showed the best results, and are slightly better than our chosen model. Lastly, training with global and local loss yielded similar results (**Figure S15A**), and so too did computing the loss on both the rows and columns compared to only considering the rows (**Figure S15 B**).

## Embedding Visualizations

To create the t-SNE plots in **Figures 2, 3, 5, S2-S4**, and **S7-S10** and in the CLSS viewer, we used 1000 iterations and a perplexity value of 30. We visualized the probability density function of the distances between embeddings using the Gaussian kernel density estimator (gaussian_kde routines) from the scipy.stats package.

## Code Availability

Our codebase (including training code) and the model weights are available on our GitHub (https://github.com/guyyanai/CLSS).

## Acknowledgements

This work was supported by Israel Science Foundation (ISF) grant 1764/21, and by the Minerva Foundation. N.B.T. is supported by the Abraham E. Kazan Chair in Structural Biology, Tel Aviv University. G.A. was supported in part by a fellowship from the Edmond J. Safra Center for Bioinformatics at Tel-Aviv University. L.M.L acknowledges support from Human Frontiers Science Program (HFSP) grant RGEC29/2025.

