## Supplementary Text & Figures for "Contrastive learning unites sequence and structure in a global representation of protein space"

### Supplementary figures

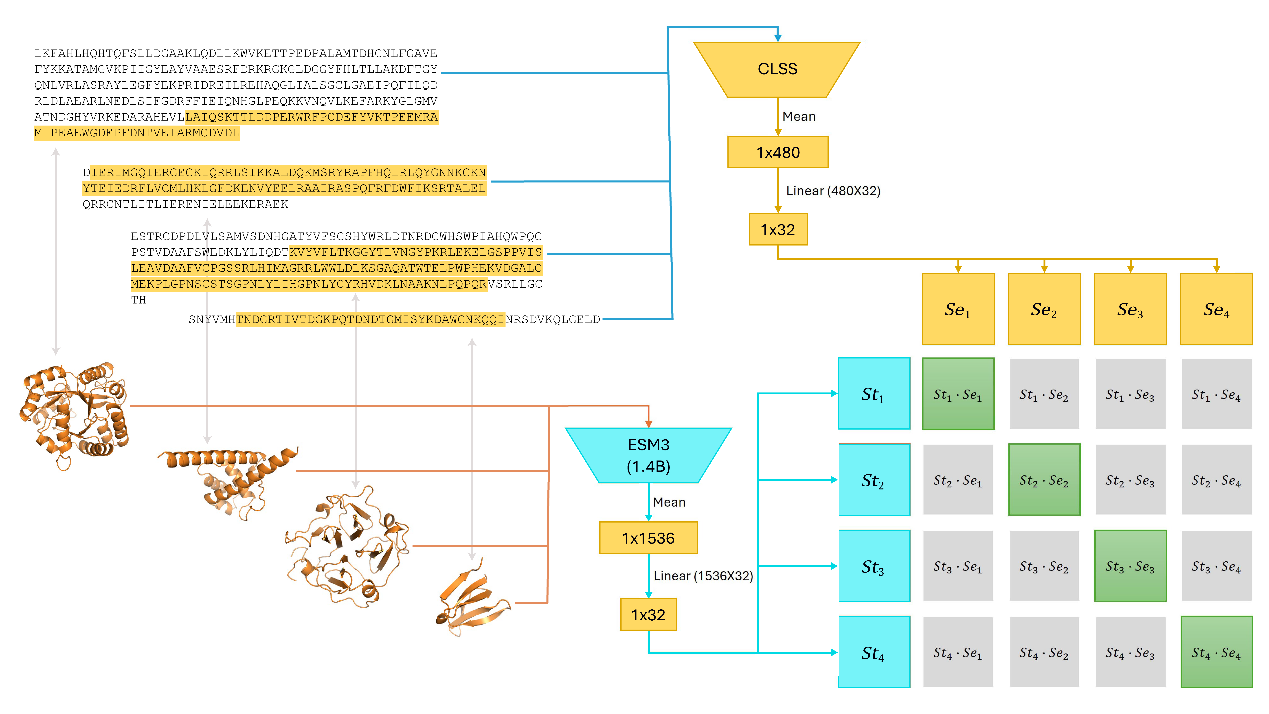

Figure S1: An illustration of CLSS: our CLIP-inspired contrastive model's architecture and training procedure. The training updates the weights in the sequence tower (ESM2 architecture and initial weights) and its adapter network, and the adapter network of the structure tower (ESM3 architecture and weights). During our training process, we calculate the embeddings using both towers: for a sampled sub-sequence (light blue), and the structure (orange), and maximize the dot-product similarities between embeddings corresponding to the same domain (marked in green), while minimizing other dot-product similarities (gray), using cross-entropy loss.

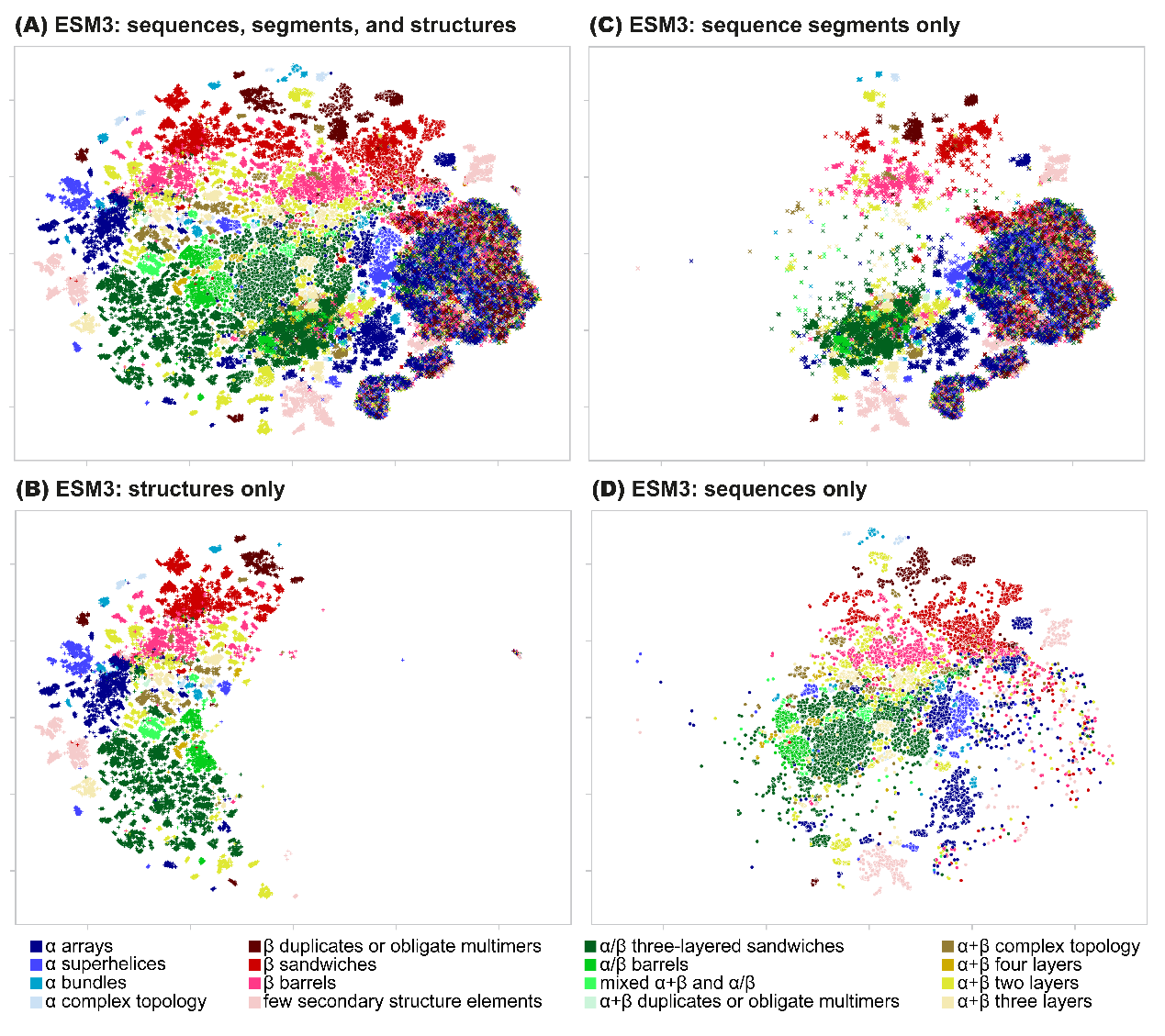
Figure S2: ESM3 maps of ECOD domains (Dataset 1) – a larger version of Figure 3, panels 2 (A-D) in the main manuscript. For each domain, we calculate the ESM3 embeddings by the structure, sequence, and a random sequence segment. Each point represents one of the modalities of a domain colored according to the label of its ECOD architecture. In this enlarged view we see that the structure modality is best aligned with the ECOD labels, the sequence modality less so, and the signal of the embeddings of the random sequence segment is very weak.

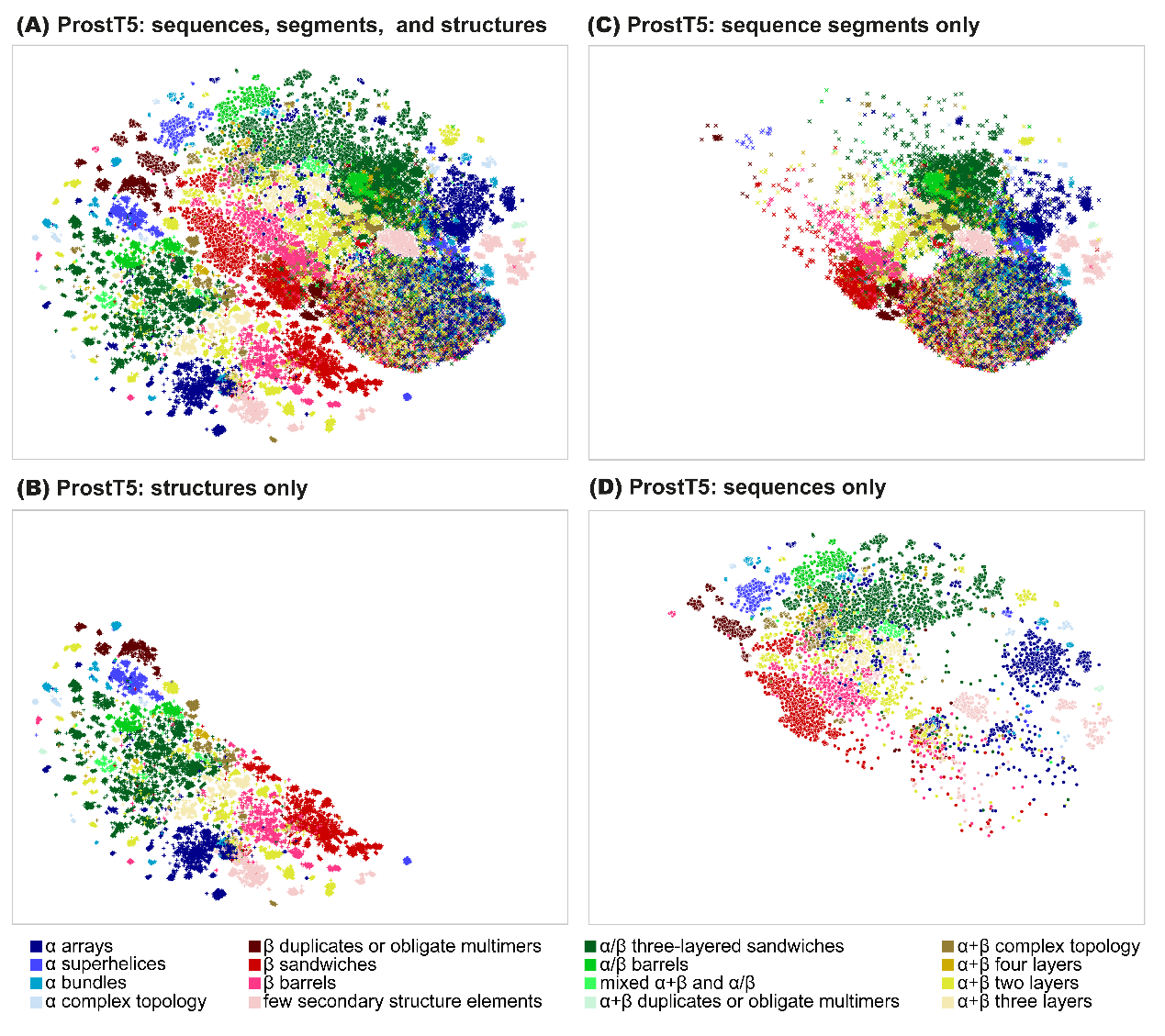
Figure S3: ProstT5 maps of ECOD domains (Dataset 1) – a larger version of Figure 3, panels 3 (A-D) in the main manuscript. For each domain, we calculate the ProstT5 embeddings by the structure, sequence, and a random sequence segment. Each point represents one of the modalities of a domain colored according to the label of its ECOD architecture. In this enlarged view we see that both the structure modality and sequence modality capture the ECOD labels well, and the random sequence segment modality far less so.

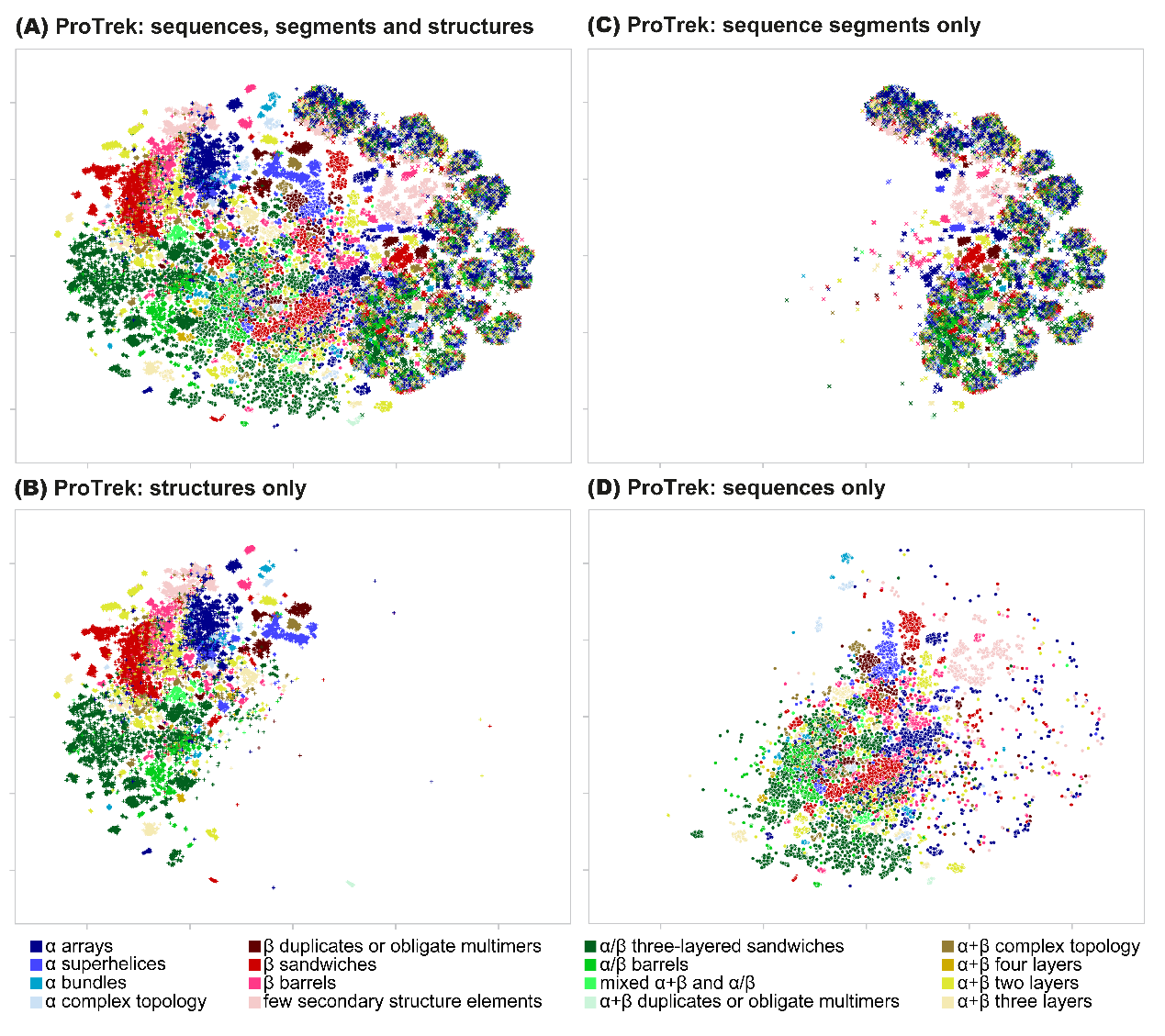
Figure S4: ProTrek maps of ECOD domains (Dataset 1) – a larger version of Figure 3, panels 4 (A-D) in the main manuscript. For each domain, we calculate the ProTrek embeddings by the structure, sequence, and a random sequence segment. Each point represents one of the modalities of a domain colored according to the label of its ECOD architecture. In this enlarged view we see that the agreement of the structure modality with the ECOD labels is the strongest, far less so for the sequence modality, and when considering the random sequence segment modality there is virtually no signal.

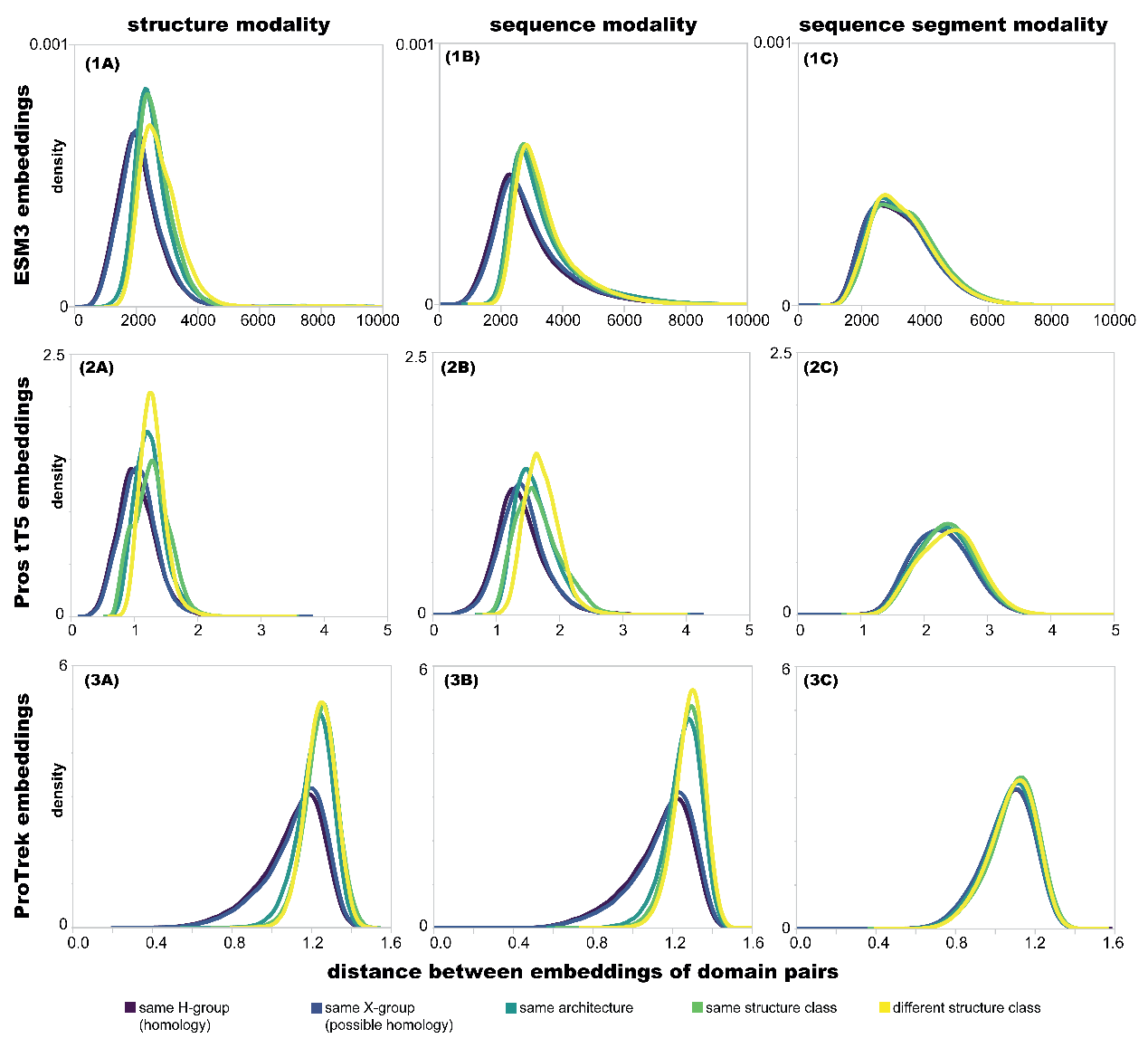

Figure S5: Estimated distributions of distances between pairs of ESM3, ProstT5, and ProTrek embeddings for domains in Dataset 1, a comparison with respect to Figure 4. The distributions of distances of domain pairs with the same ECOD label at different levels of the hierarchy. Distance distributions for pairs with the same H-group label are shown in purple through pairs with different structure class in yellow. For the sequence and structure modalities, there is a separation between the distributions of pairs with the same H-group label or X-group labels, from the pairs that are further away. Unlike CLSS, the separation between all 5 distributions is less pronounced. As for the segment modality, the distributions of the different sets of domain pairs are virtually the same, showing that there is no signal that agrees with our expectation from the human curated knowledge of the ECOD hierarchy. We do not show the distributions of high distances between different modalities of the same domain for ESM3, ProstT5, and ProTrek embeddings, as it is evident from their maps that these occupy different regions of space.

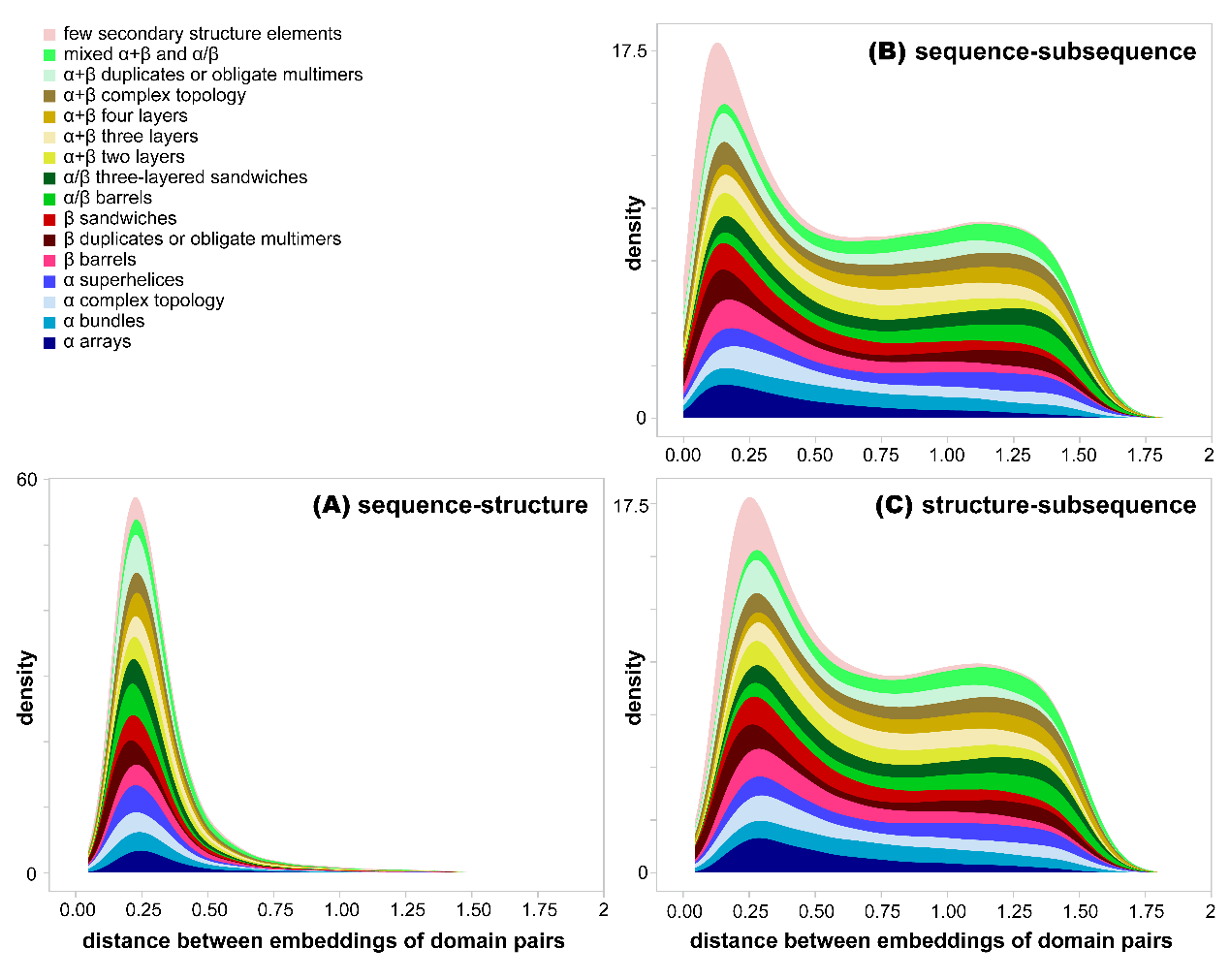

Figure S6: Distributions of the distances between CLSS embeddings of different modalities for domains in dataset 1 (an extension of the insert in Figure 4). The distributions are grouped and colored by ECOD architecture.

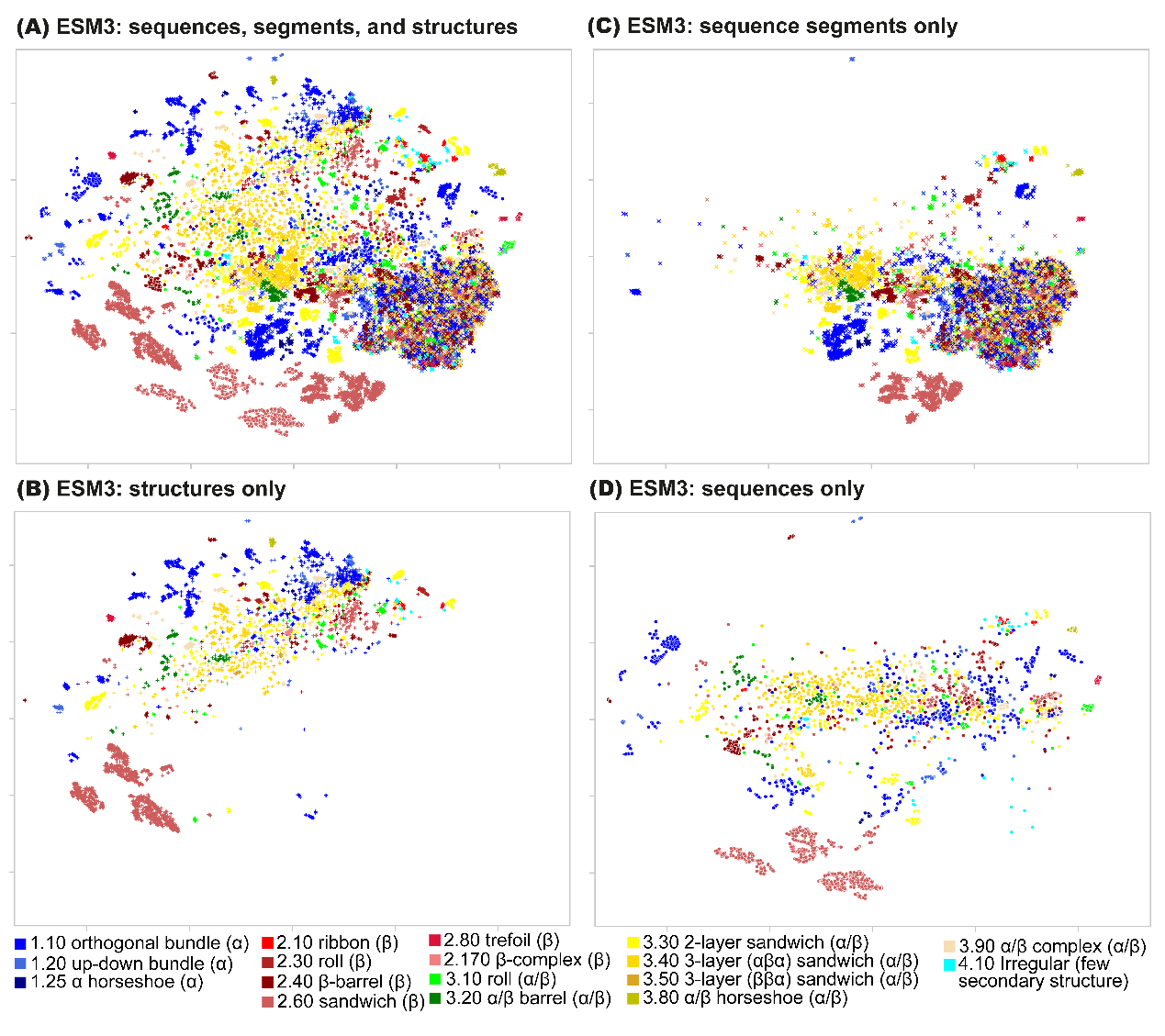
Figure S7: Maps based on ESM3 embeddings of CATH domains with similar sequences and topologically different structures (Dataset 2). These maps are in comparison to the CLSS maps for this dataset shown in Figure 5. For each domain, we consider three modalities: sequence, structure, and a random sequence segment, and compute a t-SNE projection of the embeddings. Each point represents one of the modalities of one domain colored according to the label of its CATH architecture. (A) All three modalities. Sequences are marked by circles, structures by ‘+’, and random sequence segments by ‘x’. (B) Structure embeddings. (C) sequence segment embeddings. (D) Sequence embeddings. Even though the sequence and structure modalities are not co-embedded near each other, ESM3 embeddings of these metamorphic proteins fail to successfully group the proteins according to their CATH architecture in both these modalities. In the sequence sub-segment modality, the signal is generally lost.

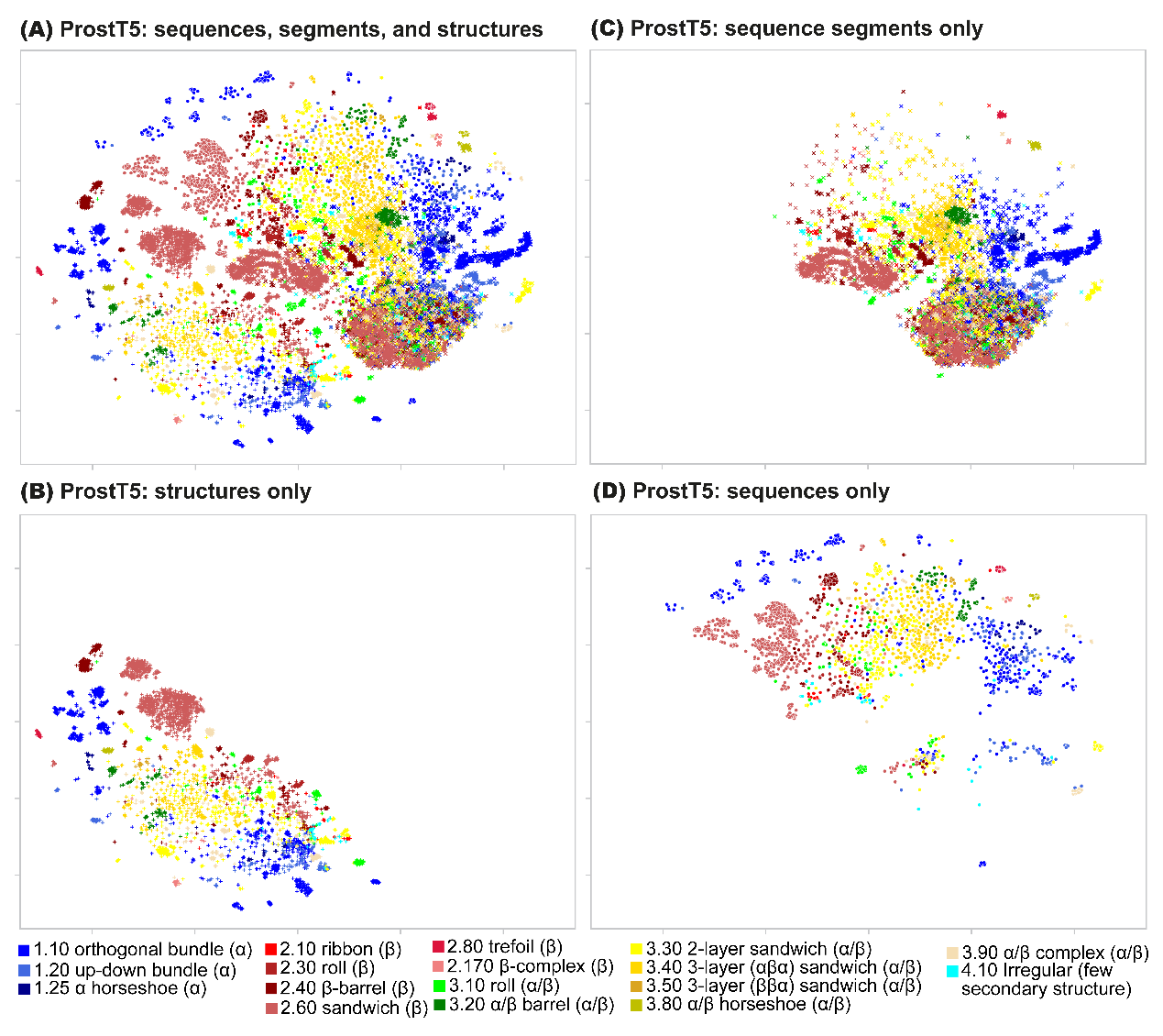
Figure S8: Maps based on ProstT5 embeddings of CATH domains with similar sequences and topologically different structures (Dataset 2). These maps are in comparison to the CLSS maps for this dataset shown in Figure 5. For each domain, we consider three modalities: sequence, structure, and a random sequence segment, and compute a t-SNE projection of the embeddings. Each point represents one of the modalities of one domain, colored according to the label of its CATH architecture. (A) All three modalities. Sequences are marked by circles, structures by ‘+’, and random sequence segments by ‘x’. (B) Structure embeddings. (C) sequence segment embeddings. (D) Sequence embeddings. ProstT5 successfully groups the proteins according to their CATH architecture in the structure and sequence modalities, but in the sequence sub-segment modality, the signal is generally lost.

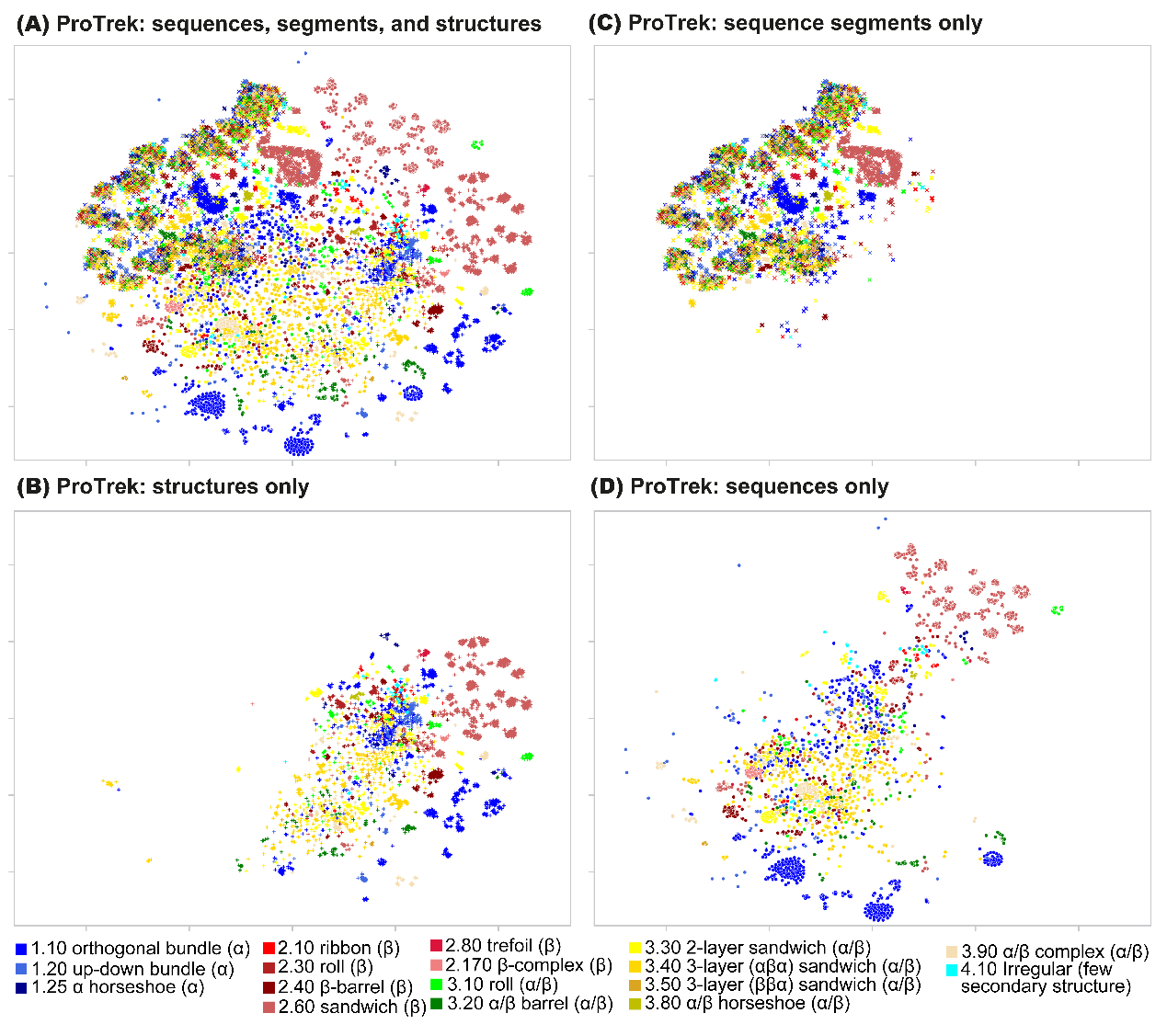

Figure S9: Maps based on ProTrek embeddings of CATH domains with similar sequences and topologically different structures (Dataset 2). These maps are in comparison to the CLSS maps for this dataset shown in Figure 5. For each domain, we consider three modalities: sequence, structure, and a random sequence segment and compute a t-SNE projection of the embeddings. Each point represents one of the modalities of one domain colored according to the label of its CATH architecture. (A) All three modalities. Sequences are marked by circles, structures by ‘+’, and random sequence segments by ‘x’. (B) Structure embeddings. (C) sequence segment embeddings. (D) Sequence embeddings. ProTrek embeddings of these metamorphic domains correlate to some extent with CATH architecture annotations in the structure, and less so in the sequence modality. In the sequence sub-segment modality, the signal is generally lost.

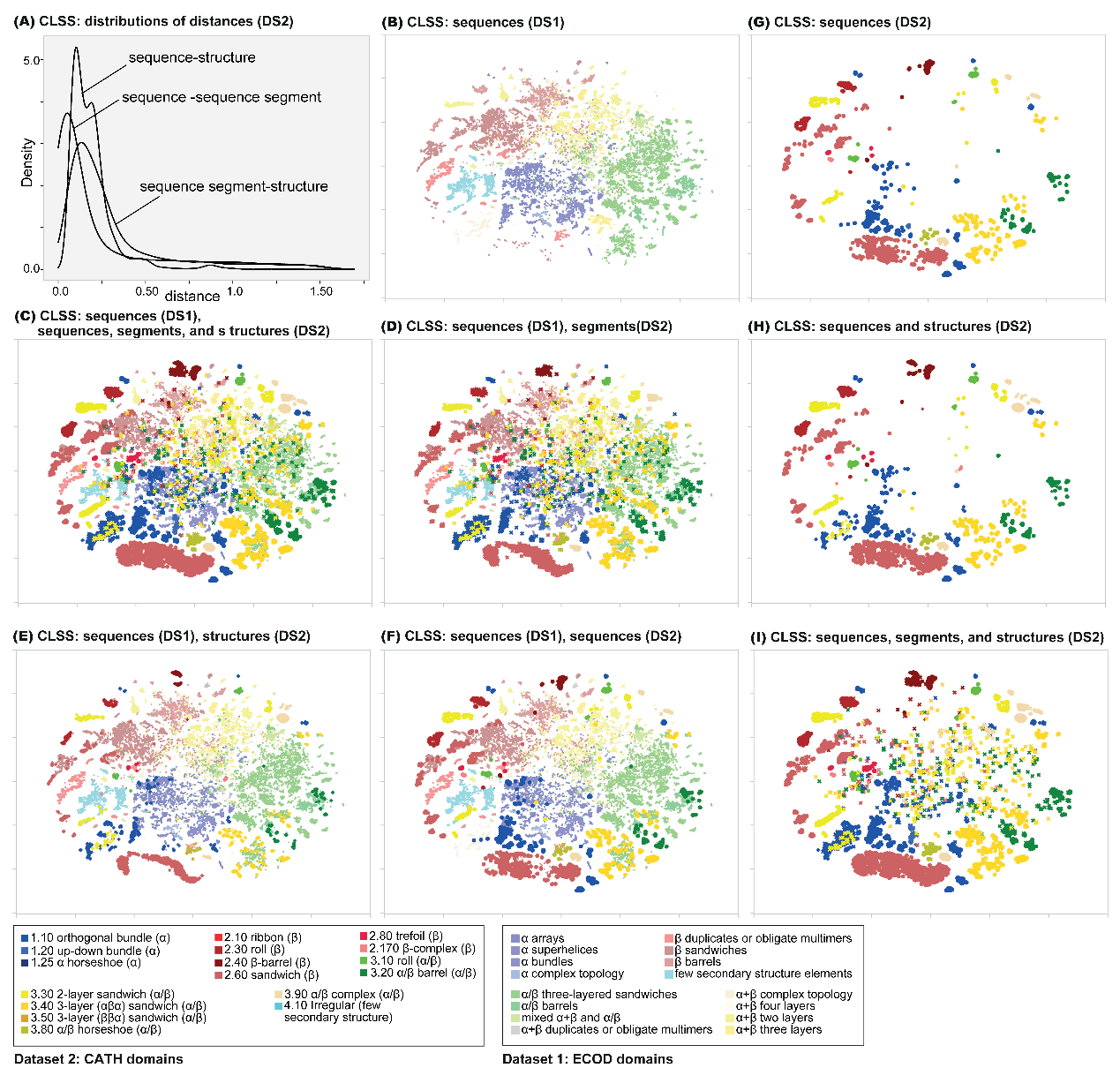

Figure S10: Further characterization of CLSS embeddings of Dataset 2 (DS 2) of metamorphic proteins, when mapped together with the ECOD domains in Dataset 1 (DS 1). (A) The distribution of distances between two modalities of the same domain in the Dataset 2. In all cases the distances are relatively small (comparable to those shown in Figure 3A for Dataset 1). To place the embeddings of Dataset 2 within the context of protein structure space, we calculated the t-SNE projection of the sequence embeddings of the ECOD domains and the three modalities of Dataset 2. The embeddings are colored by architecture, in Dataset 2 using vivid colors, and in Dataset 1 using muted colors. The different panels show different subsets of these embeddings. (B) The ECOD domains of Dataset 1 map, shown for context, (C-F) The ECOD domains along with all three modalities, and each of them separately. (G-I) Only the embeddings of Dataset 2 of metamorphic proteins: (G) only sequence modality, (H) the sequence and structure modalities, (I) all three modalities. Panels (G-H) show that the embeddings of the sequence and structure modalities generally fall in the periphery of the protein sequence space marked by the embeddings of the ECOD domains in Dataset 1.

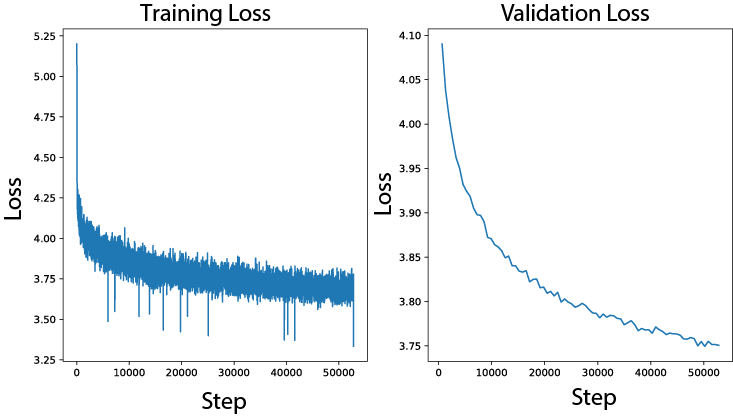

Figure S11: Training and validation loss graphs of the chosen architecture of CLSS, with an adapter of size 32, that was trained for 80 epochs on subsequences with a minimum length of 10.

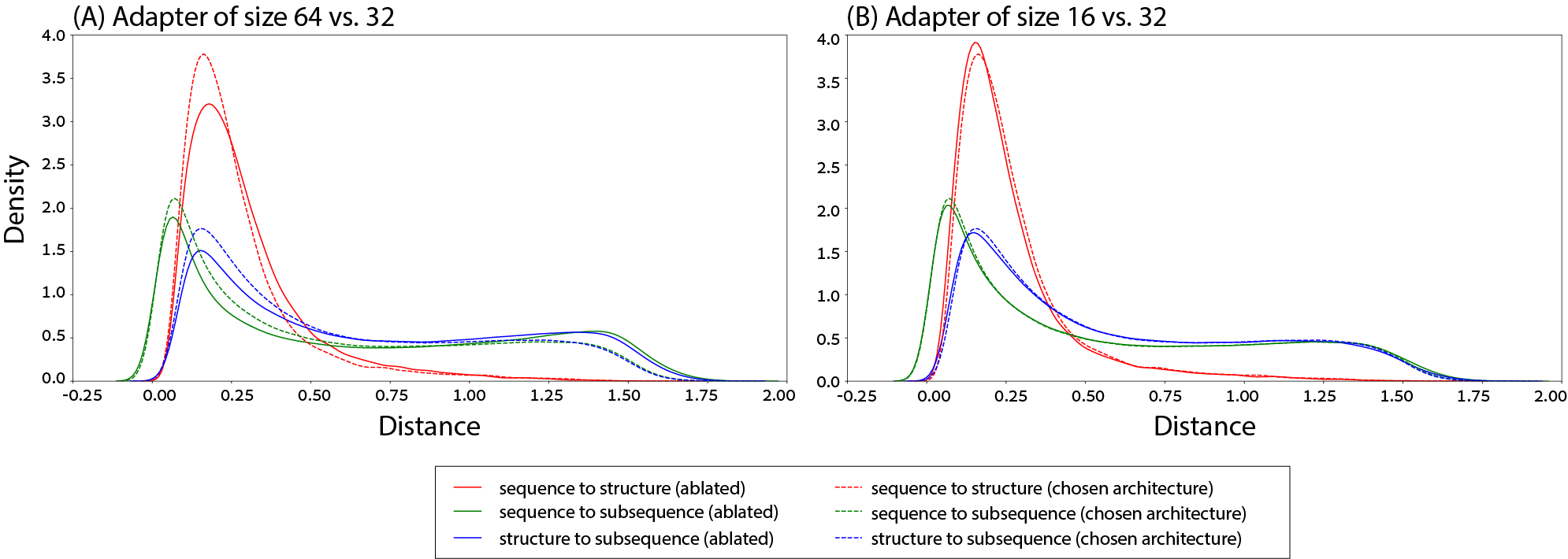

Figure S12: Distributions of distances between the three different modalities of each domain in the validation set computed by different models. We compare our chosen architecture that has adapters of size 32, to alternate architectures with adapters of size 16 (panel A), and 64 (panel B). Panel A shows the chosen architecture outperforming the alternative across the board, while panel B shows very similar results between the chosen architecture and the alternative.

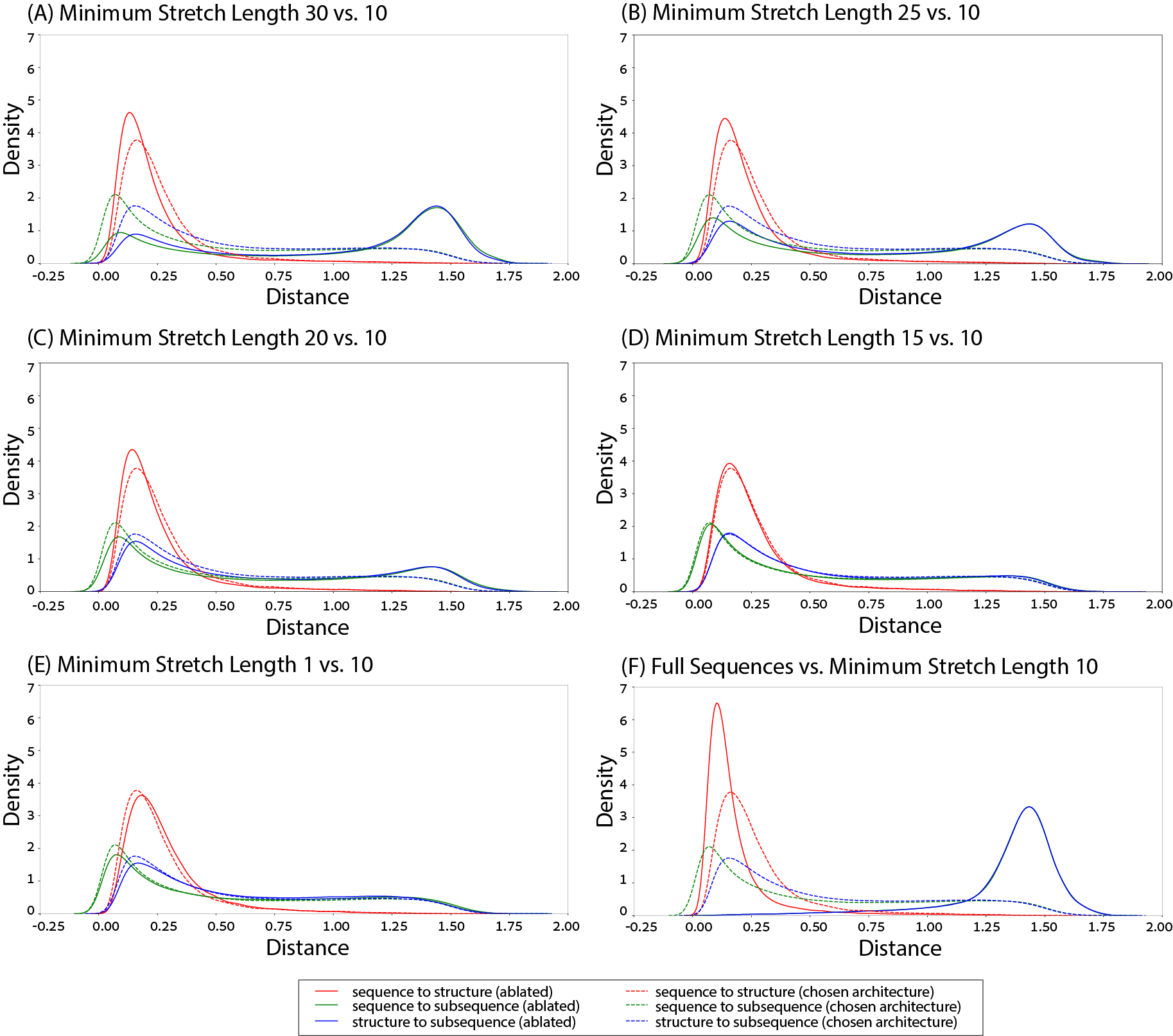

Figure S13: Distributions of distances between the three different modalities of each domain in the validation set computed by different models. We compare our chosen architecture that trained on subsequences with a minimum length of 10 to models trained on subsequences with a minimum length of 30 (panel A), 25 (panel B), 20 (panel C), 15 (panel D), 1 (panel E) and an additional model trained on full sequences (panel F). Panels A-C show our chosen architecture vastly outperforming the alternatives in distances between subsequences and sequences/structures, while having higher distances between sequences and structures than the alternative architectures. Panel D shows similar results between the chosen architecture to the ablated one. Panel E shows the chosen architecture slightly outperforming the ablated architecture. Panel F shows the chosen architecture performing worse on distances between sequences and structures, while performing substantially better on distances involving subsequences.

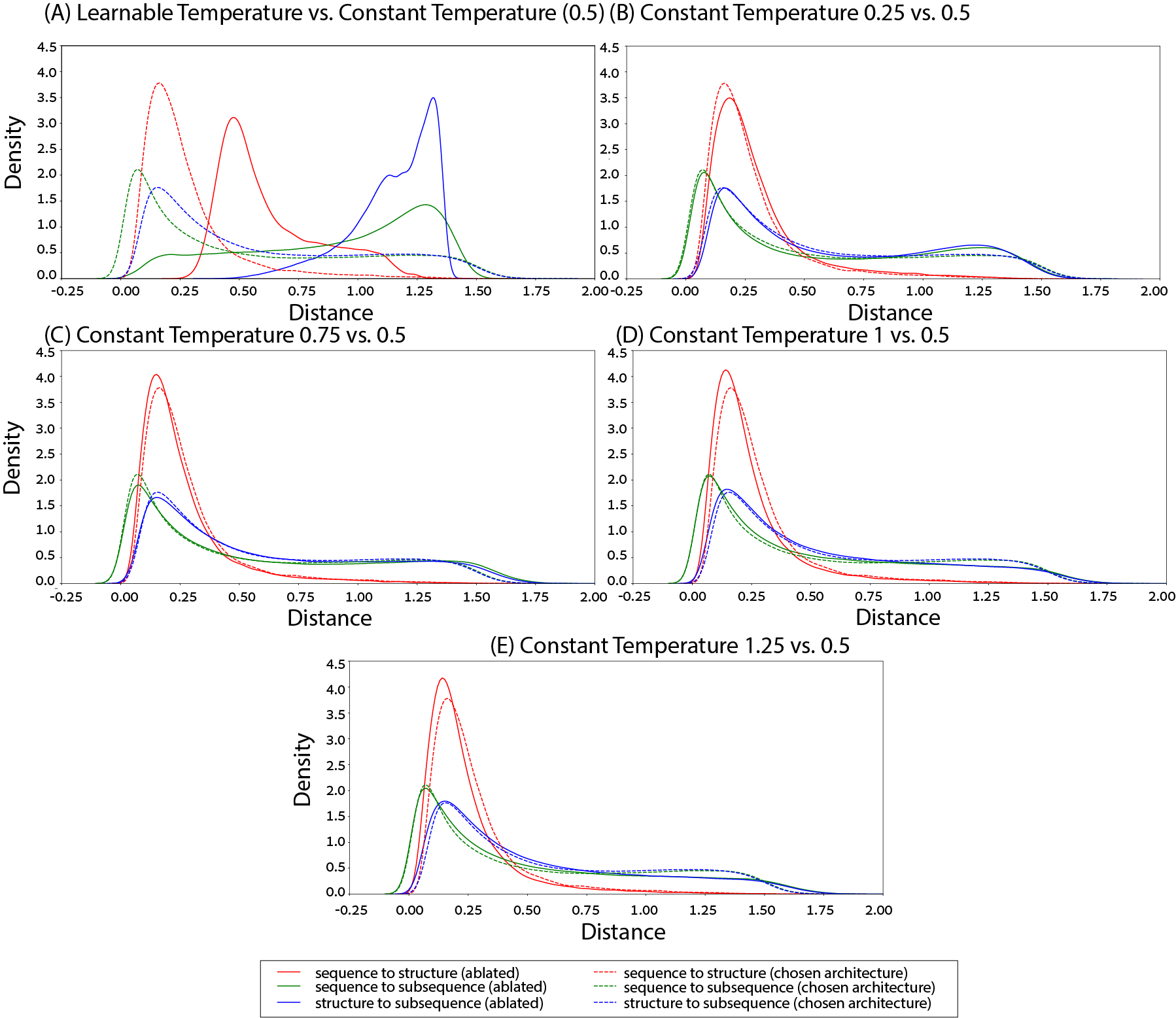

Figure S14: Distributions of distances between the three different modalities of each domain in the validation set computed by different models. We compare our chosen architecture that trained using a constant temperature of 0.5 to models trained using a learnable temperature (panel A), or constant temperatures of 0.25 (panel B), 0.75 (panel C), 1 (panel D), 1.25 (panel E). Panel A shows the chosen architecture performing significantly better than the ablated model. Panel B shows the chosen model performing slightly better than the alternative. Panel C shows mixed results, and panels D,E show the ablated models examine a slight improvement over the chosen architecture.

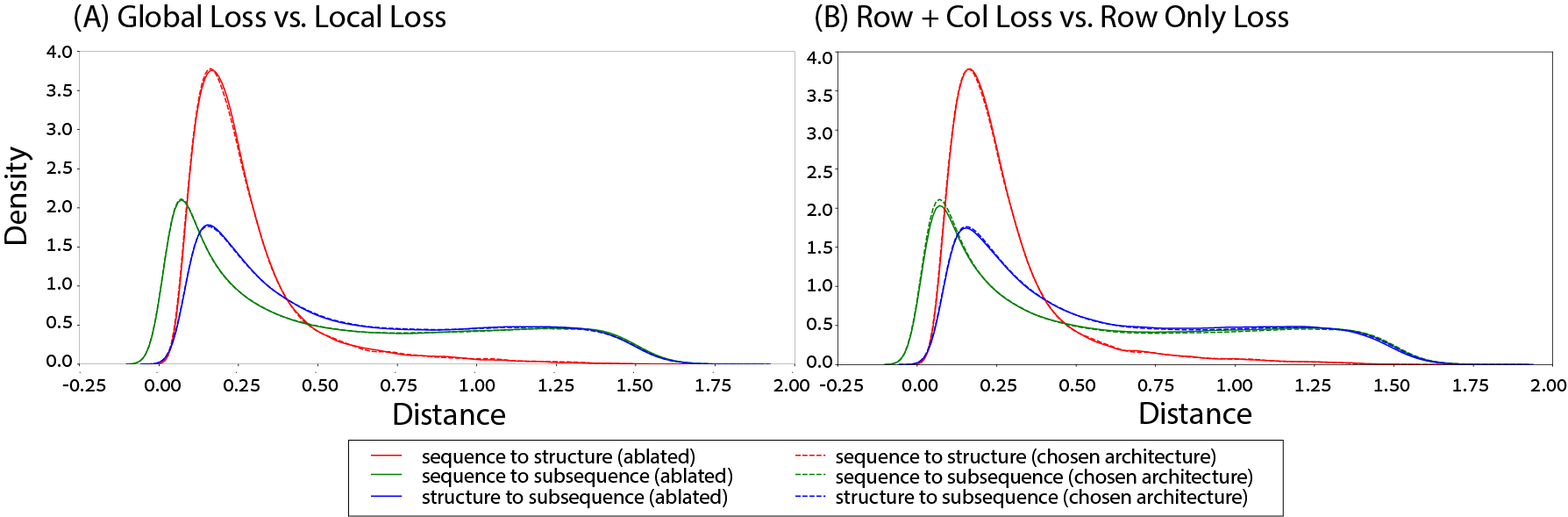

Figure S15: Distributions of distances between the three different modalities of each domain in the validation set computed by different models. We compare our chosen architecture that trained using local, row-only loss, to a model that trained on global, row-only loss (panel A), and a model trained on local, row-and-column loss (similar to the CLIP loss) (panel B). Both panels show very similar results with a slight edge to the chosen architecture.

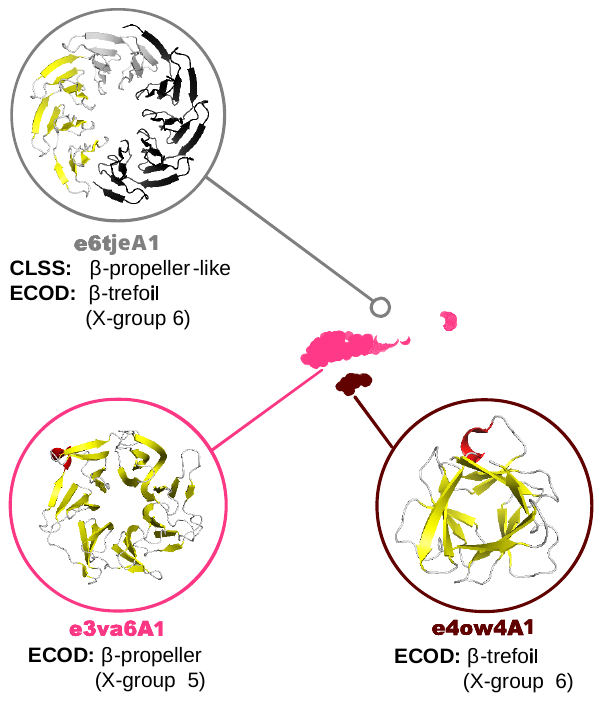

Figure S16: Another example comparison between the ECOD hierarchy and CLSS maps highlighting the challenge of automatic classification of a designed protein with sequence-based HMMs. In ECOD version 291, e6tjeA1 is part of a designed β-propeller with β-trefoil-like sequence characteristics. See **Supplemental Text** for more details.

### Supplementary text

We show another example that highlights the challenges facing sequence-based HMMs when used for classification. We have previously observed sequence and structure connections between the β-trefoils (X-group 6) and β-propellers (X-group 5) [1,2]. Consistent with these results, the CLSS embeddings of these two β-folds form adjacent but distinct clusters. The domain e6tjeA1 annotated by the automatic ECOD pipeline as a β-trefoil, clusters with the β-propellers in the CLSS embedding space across all three modalities (**Figure S16**). Inspection of the structure reveals that 6tje is a Rosetta-designed 9-bladed β-propeller called Cake5 [3], consistent with the CLSS embedding. The classification of e6tjeA1 by the automatic ECOD pipeline (version 291) as a β-trefoil suggests that these folds are indeed evolutionarily related and provides an instructive example: Briefly, Cake5 must dimerize to form a complete β-propeller and the ECOD pipeline divided one chain of Cake5 into two separate domains: The first “β-trefoil domain” (e6tjeA1) comprises 3 β-propeller blades with a total of 12 β-strands, the same number as in a canonical β-trefoil. The second “domain” (e6tjeA2) encompasses just 7 β-strands and is correctly classified by ECOD as a β-propeller, despite being a smaller segment. As Cake 5 is a designed sequence (which can differ significantly from known protein families of the same fold), a repetitive sequence (where the number of repeats can vary between related domains, complicating the identification of domain boundaries), and a homo-oligomer (where the full domain is split across two chains), it is an exceptionally difficult protein to classify automatically.
